# The m^6^A landscape of polyadenylated nuclear (PAN) RNA and its related methylome in the context of KSHV replication

**DOI:** 10.1101/2021.04.02.438257

**Authors:** Sarah Elizabeth Martin, Huachen Gan, Gabriela Toomer, Nikitha Sridhar, Joanna Sztuba-Solinska

## Abstract

Polyadenylated nuclear (PAN) RNA is a non-coding transcript involved in Kaposi’s sarcoma-associated herpesvirus (KSHV) lytic reactivation and regulation of cellular and viral gene expression. We have shown that PAN RNA has a dynamic secondary structure and protein binding profiles that can be influenced by the epitranscriptomic modifications. N^6^-methyladenosine (m^6^A) is an abundant signature found in viral and virus-encoded RNAs. Here, we combined an antibody-independent next-generation mapping with direct RNA sequencing to elucidate the m^6^A landscape of PAN RNA during the KSHV latent and lytic stages of infection. Using a newly developed method, termed <u>S</u>elenium-modified deoxythymidine triphosphate reverse transcription and <u>L</u>igation <u>A</u>ssisted <u>P</u>CR analysis of m^6^A (SLAP), we gained insight into the fraction of modification at identified sites. Using comprehensive proteomic approaches, we identified writers, erasers, and readers that regulate the m^6^A status of PAN. We verified the temporal and spatial subcellular availability of the methylome components for PAN modification by performing confocal microscopy analysis. Additionally, the RNA biochemical probing outlined structural alterations invoked by m^6^A in the context of full-length PAN RNA. This work represents the first comprehensive overview of the dynamic interplay between the cellular epitranscriptomic machinery and a specific viral RNA.

## INTRODUCTION

Kaposi’s sarcoma-associated herpesvirus (KSHV) is etiologically linked to several malignancies, including Kaposi’s sarcoma (KS), primary effusion lymphoma (PEL), and Multicentric Castleman’s disease (1–4). Like all herpesviruses, the KSHV infectivity cycle is bipartite and includes latent and lytic stages of infection. During the latency, KSHV persists as episomal DNA and expresses only a few of its genes. The full repertoire of viral genes expression occurs in a highly ordered manner during the KSHV lytic replication (5). Polyadenylated nuclear (PAN) RNA is a multifunctional long non-coding (lnc) RNA produced at high levels during KSHV lytic reactivation (6), although it is also expressed during KSHV latency in cultured cells (7). PAN RNA regulates cellular and viral gene expression by recruiting chromatin-modifying complexes to cellular and viral DNA (8–10). Another study showed that PAN facilitates late viral mRNA export to the cytoplasm, increasing the levels of viral mRNAs available for the expression of late lytic products (11). PAN also acts as a scaffold for viral proteins ORF50 and ORF57, and RNA polymerase II at chromatin, leading to the execution of the viral lytic program (8). We have previously shown that PAN RNA has dynamic secondary structure and protein binding profiles, which change depending upon the biological context (12). This work constitutes the most extensive structural characterization of viral lncRNA and its interactome inside the living cells and virions, providing a broad framework for understanding PAN’s roles in KSHV infection.

Recent studies have shown that the structure (13–18), metabolism (19–21), and function (18, 22–24) of coding and non-coding RNAs can be modulated by over 100 different types of chemical modifications, that are collectively referred to as the epitranscriptome. N^6^-methyladenosine (m^6^A) is the most abundant of those, as it has been found in over 25% of transcripts expressed in mammalian cells (25), as well as in viral RNA genomes (26, 27) and virus-encoded RNAs (28, 29). m^6^A has been shown to impact nearly every aspect of RNA biology, from splicing to translation, and all the way through stability and decay (30–32). The breadth of m^6^A impact has been attributed to creation of new sites for protein recognition, in part via local structural changes to the modified RNA (15, 33, 34). However, the biological significance of m^6^A, especially in the context of a specific viral transcript, has not been defined.

Previous studies have outlined the mechanism regulating the m^6^A status of KSHV coding RNAs (25, 35–38). It has been shown that the modification is mediated by a complex of methyltransferases (METTL3/14) and Wilms tumor 1 associated protein (WTAP), which are recruited to target RNA via RNA-binding motif 15 protein (RBM15) (39). The fat mass and obesity-associated protein (FTO) reverses this process and acts as m^6^A eraser (36, 41, 44). Additionally, the biological functions of m^6^A can be mediated by reader proteins (38). For example, heterogeneous nuclear ribonucleoprotein C (HNRNPC), responsible for RNA splicing and stability, selectively binds m^6^A, where the modification acts as a ‘switch’, destabilizing an RNA helix and exposing a single-stranded HNRNPC binding motif (33). The structural influence of this ‘m^6^A-switch’ also dictates the binding of YTH domain-containing proteins. In the context of KSHV infected cells, the YTHDC1 binding potentiates viral mRNAs splicing (36), while YTHDF2 overexpression facilitates the degradation of KSHV coding transcripts (42). In addition, Staphylococcal nuclease domain-containing protein 1 (SND1) has been reported to bind and stabilize the KSHV ORF50 mRNA, supporting viral replication (37). Nonetheless, the complexity of the KSHV transcriptome and the likely related epitranscriptome, expands beyond coding RNAs and includes many non-coding transcripts, with PAN leading in abundance and significance for KSHV replication (5, 43, 44). As of today, the m^6^A landscape and its dynamics for any of these viral non-coding transcripts, have not been elucidated.

In this study, we applied an antibody-independent next-generation sequencing-based approach and direct RNA sequencing to tackle the PAN m^6^A landscape during the latent and lytic stages of KSHV replication. We developed a novel approach, <u>S</u>edTTP-RT and <u>L</u>igation <u>A</u>ssisted <u>P</u>CR analysis (SLAP), to quantify the modified fraction, which is a critical parameter for investigating m^6^A biological significance. Using target-specific RNA crosslinking, affinity capture and mass-spectrometry, we delineated the cellular methylome that guides m^6^A installation on PAN RNA and potentially conveys its phenotypic effect. Furthermore, confocal microscopy analyses identified the temporal and spatial subcellular availability of these components for PAN m^6^A modification. We also addressed the influence of m^6^A on the local and global secondary structure of full-length PAN RNA by performing biochemical RNA structure probing (SHAPE-MaP). This study provides the first comprehensive insight into the m^6^A status of a specific viral lncRNA, revealing the sophisticated interplay that exists between this viral lncRNA and cellular epitranscriptomic machinery, and creates a paradigm for future studies in the field of host-pathogen interactions.

## MATERIAL AND METHODS

### Cell lines and culture conditions

The KSHV positive body cavity-based lymphoma cell line (BCBL-1, a generous gift from Dr. Denise Whitby, NCI Frederick) was seeded at 2 x 10^5^ cells/ml and grown in RPMI-1640 medium (ThermoFisher 11875085) supplemented with 10% fetal bovine serum (FBS), 1% Penicillin/Streptomycin and 1% L-Glutamine at 37 °C in 5% CO_2_. The induction of KSHV lytic infection was performed by treating 2 x 10^7^ BCBL-1 cells with sodium butyrate (NaB, EMD Millipore 654833) to a final concentration of 0.3 mM, and collecting cells at 8-, 24-, 48-, and 72-hours post-induction (h pi).

### In vitro RNA transcription

The first 100 nucleotides (nt) of the PAN sequence with T7 promoter was synthesized using oligonucleotides listed in Table 1 and served as a template for in vitro transcription of the unmodified transcript (negative control). The RNA with the equivalent sequence but carrying two m^6^A sites was purchased from Integrated DNA Technologies (IDT, positive control). In vitro transcription of unmodified PAN transcript was performed using the MEGAscript T7 Transcription Kit (Thermo Fisher AM13345). The 100% m^6^A modified full-length PAN transcript was synthesized using the MEGAscript T7 Transcription Kit following the manufacturer’s protocol but substituting adenosine-triphosphate with N^6^-methyladenosine-5’-triphosphate (Jena Bioscience NU-1101S). RNAs were purified using the MEGAclear Transcription Clean-Up Kit (ThermoFisher AM1908).

**Table 1.**
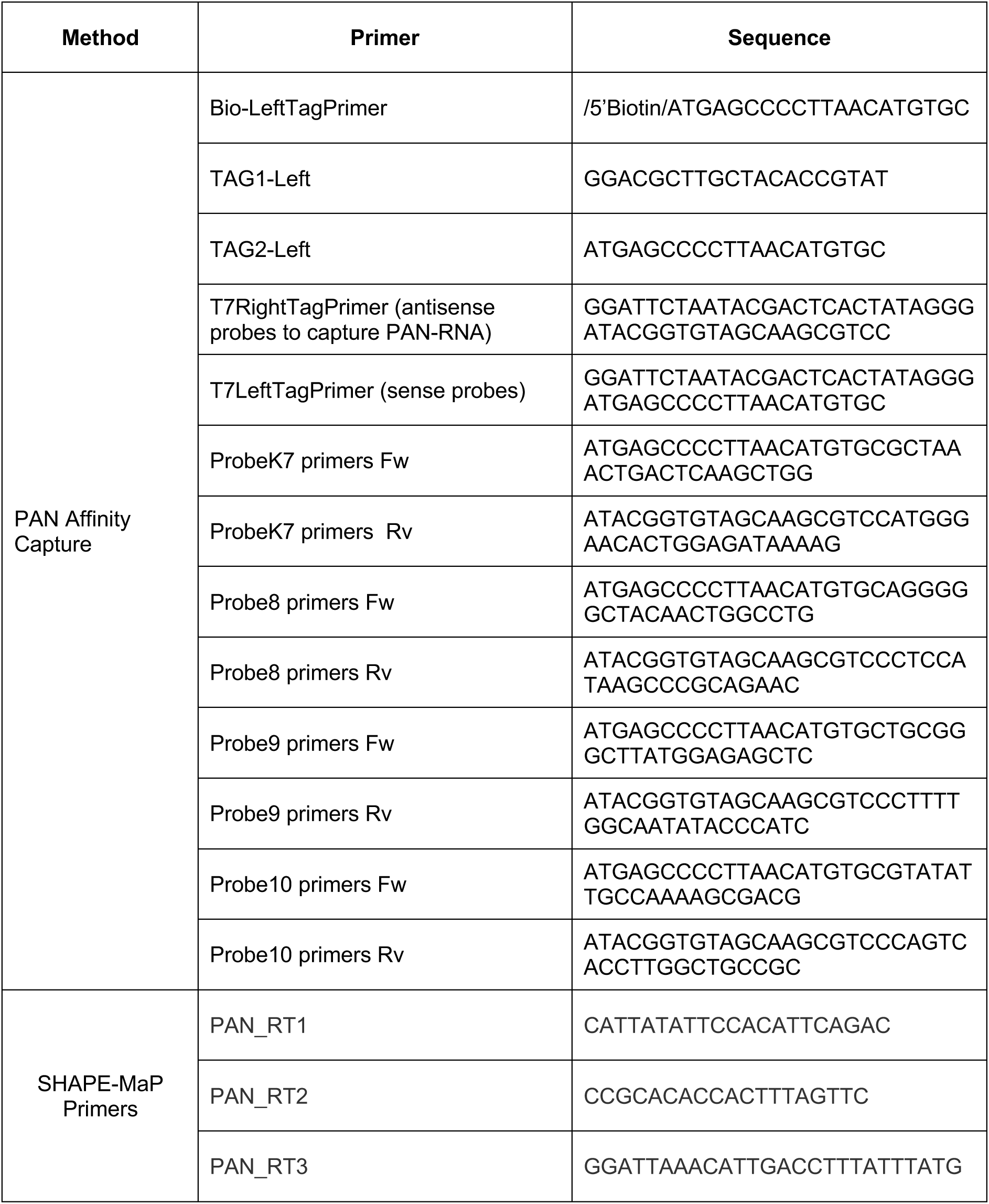
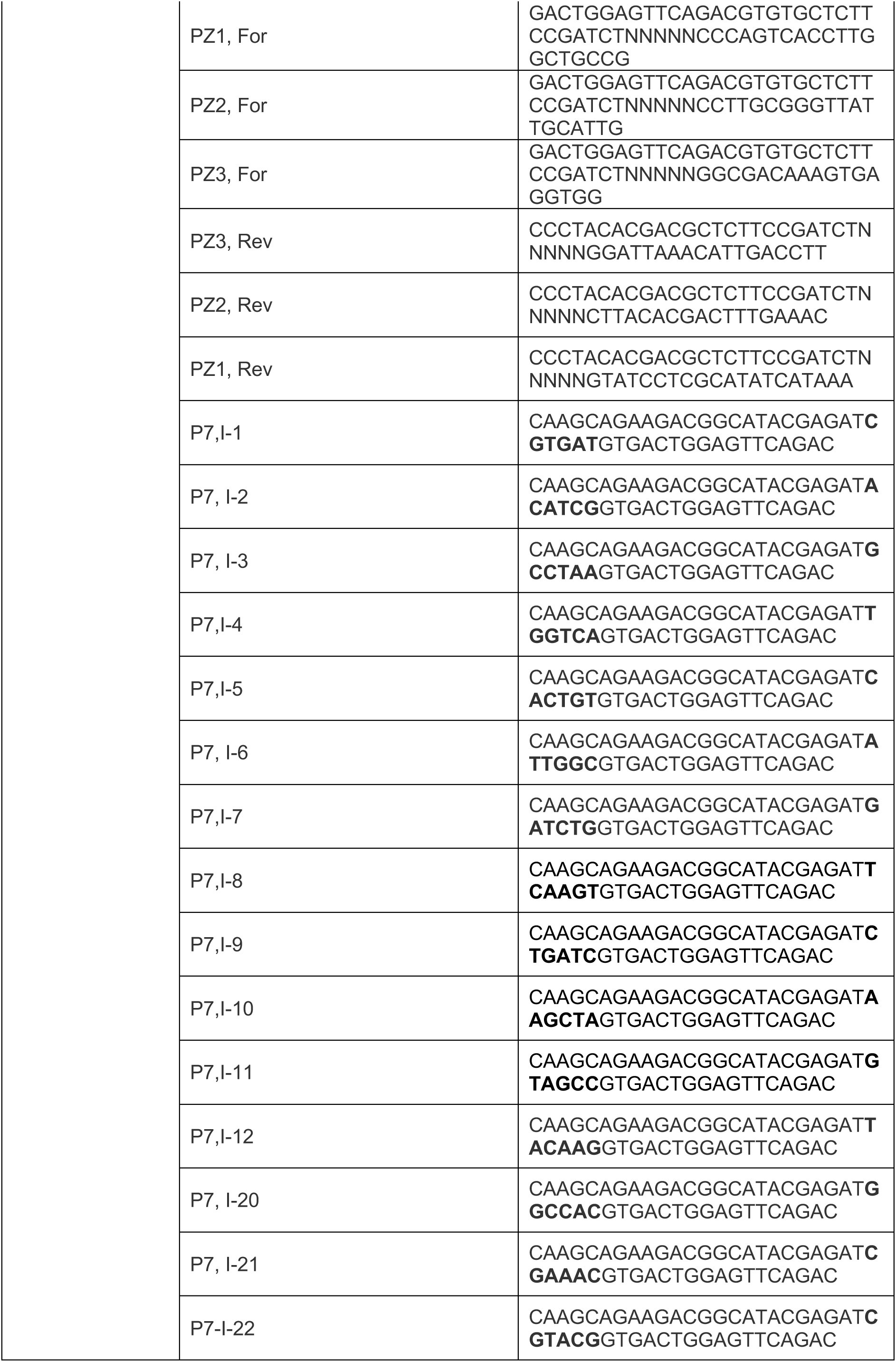
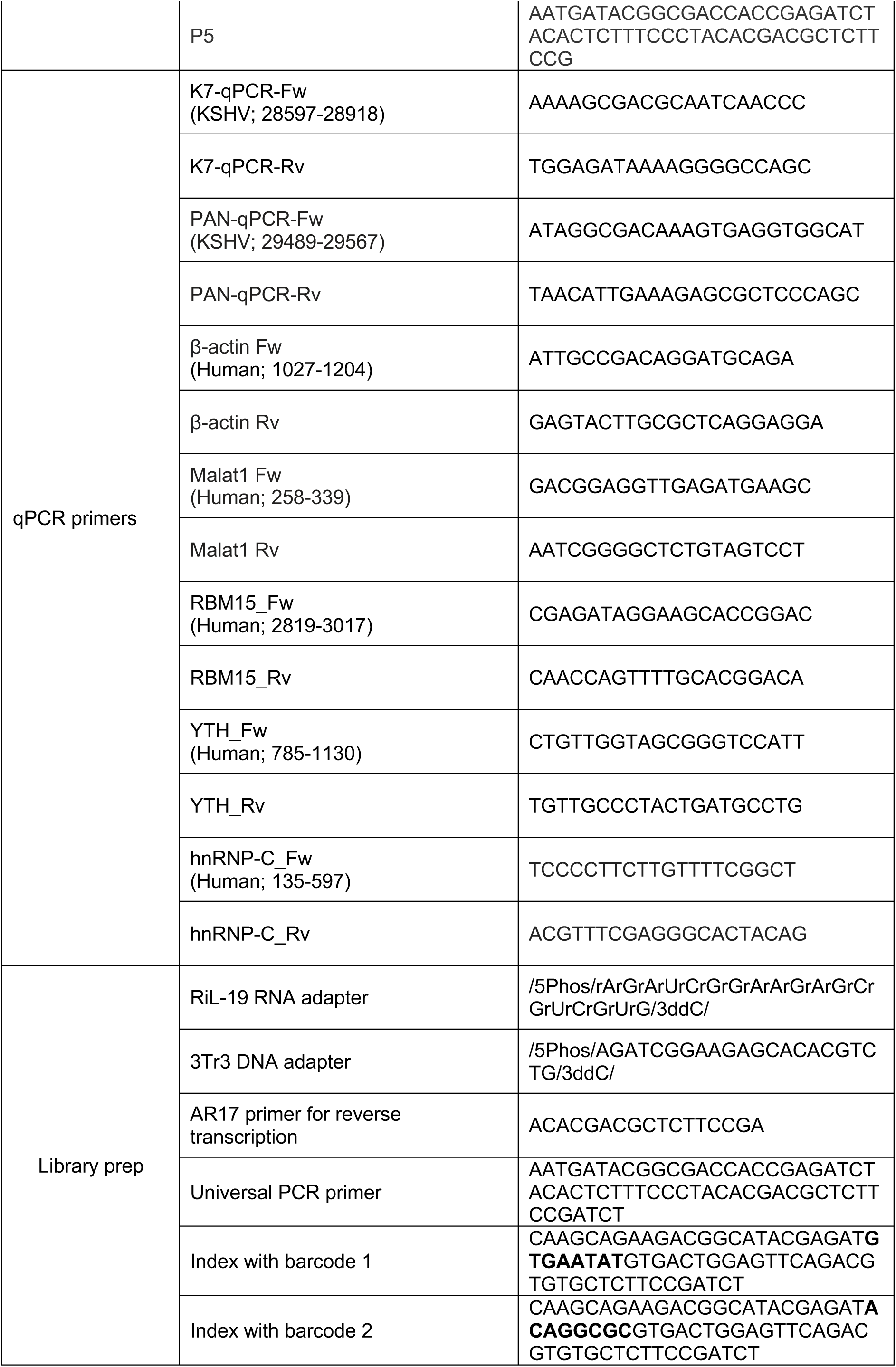
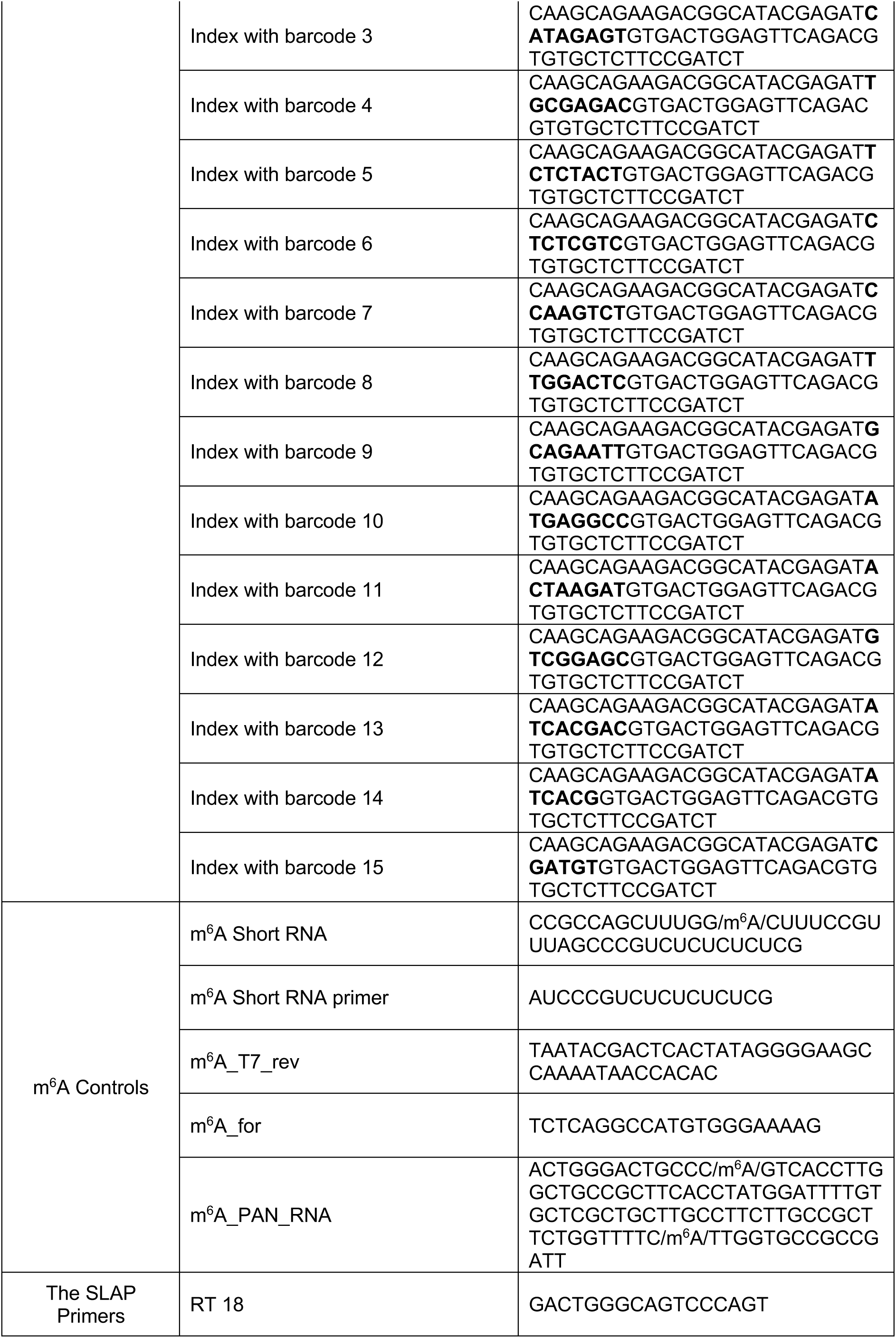
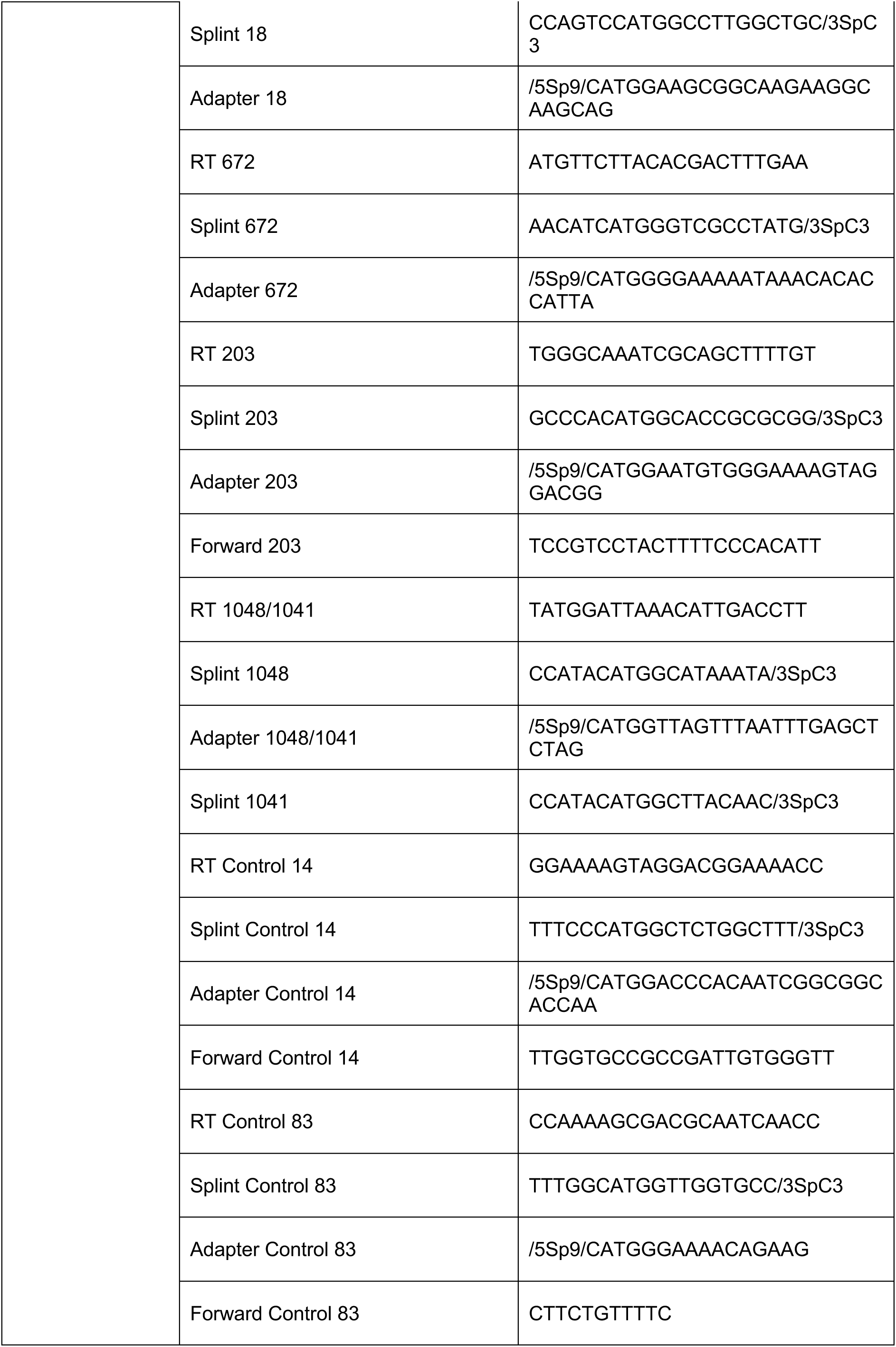
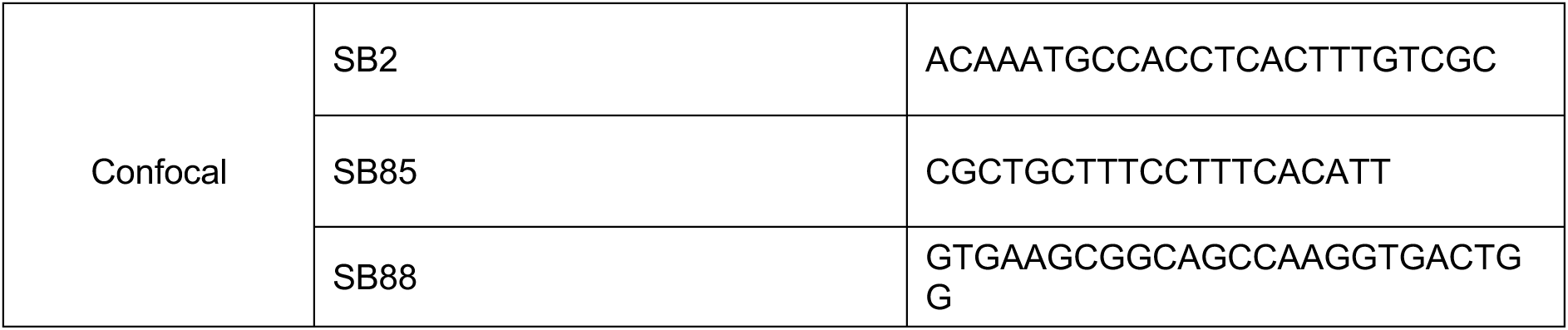
List of oligonucleotides used in this study. The following abbreviations have been used to indicate specific to the sequences: 5Biosg - 5’ biotinylation. 5Phos - 5’ phosphorylation. N - random nucleotides. 5Sp9 - triethylene glycol spacer. 3ddc - 3’ dideoxycytidine chain terminator, Fw – forward, Rv-reverse orientation. The bolded sequences indicate Illumina Tru-seq indexes.

### Probe generation

Ten biotinylated antisense probes, each 120 nt long, were designed to anneal to the PAN RNA sequence and synthesized per previously published method (18). One 45 nt long probe annealing outside the K7/PAN loci overlap was designed to perform the depletion of K7 mRNA sequence (5). Three 50 nt long probes were purchased (IDT) and used to capture human U1 RNA (positive control). DNA templates for PAN-specific probes were obtained through reverse transcription (RT) of 100 ng total RNA from the lytically-induced BCBL-1 using oligonucleotides listed in Table 1. Reverse transcription (RT) reactions (total volume of 20 µl) included 250 ng random hexamer primers (IDT 51-01-18-26), 10 mM dNTP, 1X First-strand buffer, 0.1 M DTT, 2 μl RNase OUT (40U final, ThermoFisher 10777-019), and 1 μl of SuperScript II RT (200U final, ThermoFisher 18064014). The left and right tag sequences were added during PCR amplification (Table 1). The products were purified using the PureLink PCR purification kit (ThermoFisher K310002) and used as templates for a second round of amplification to add the T7 promoter sequence in the antisense orientation. DNA products were purified and used for in vitro transcription. 10 µg of resulting RNA was used for RT using the biotinylated tag primer (Table 1). The biotinylated probes were purified using the RNA Clean & Concentration -5 purification kit (Zymo R1013).

### PAN RNA affinity capture

Total RNA was extracted using TRIzol^TM^ (ThermoFisher 15596026), followed by DNase I treatment (ThermoFisher AM1907), and purification using RNA Clean & Concentrator-5. K7 RNA was depleted by affinity capture with a biotinylated antisense K7 oligonucleotide probe (Table 1). The remaining RNA fraction was purified using SILANE beads (ThermoFisher 37002D) and fragmented using the COVARIS 8 sonicator with the following conditions: 250 seconds at 75 peak power with 26.66% duty factor, 1000 cycles, with 20 average power at 4 °C. PAN affinity capture was performed using 5 µg of total RNA, 15 pmol pool of the antisense probes (Table 1), and streptavidin-coated magnetic beads (ThermoFisher 65002).

### Quantitative reverse transcription PCR (RT qPCR)

RT qPCR was performed using SuperScript II RT with random hexamers following the manufacturer’s protocol. A 1:10 dilution of the RT reaction was used to perform a quantitative real-time PCR (qPCR) assays using Perfecta™ SYBR® Green FastMix™ Low ROX™ (Quanta BioSciences 95071). The BestKeeper ver. 1 was used to perform the repeated pair-wise correlation and regression analysis for the level of RNA expression of housekeeping genes, i.e., β-actin, glyceraldehyde 3-phosphate dehydrogenase (GADPH), U1 small nuclear (sn) RNA, and Metastasis Associated Lung Adenocarcinoma Transcript 1 (MALAT1) lncRNA during the latent and lytic stages of KSHV replication. The analysis of MALAT1 and β-actin transcripts expression resulted in the coefficient of correlation values of R = 0.98 and R = 0.94, respectively. These genes were used for data normalization.

### 4-Selenothymidine-5’-triphosphate reverse transcription (4SedTTP RT)

2.5 µg of fragmented and affinity captured PAN RNA was used for 20 μl ligation reaction that included 20 pmol of RiL-19 oligonucleotide (Table 1), 2 μl 10X T4 Ligation Buffer, 1.8 μl 100% DMSO, 100 mM ATP, 8 μl PEG-8000, 0.3 μl of RNase Inhibitor (12U final, NEB M0307L) and 1.8 µl of T4 RNA ligase (20U final, NEB M0204S), and incubated at room temperature for 90 minutes (min). Reverse transcription reaction was initiated by annealing 20 pmol of AR-17 oligonucleotide (Table 1) to the sample and incubating the reaction for 5 min at 75°C and slowly cooling to 25°C. Next, 2 µl of 4SedTTP reaction buffer (500 mM Tris-HCl pH 8.0, 500 mM KCl, 50 mM MgCl_2_, 100 mM DTT), 2 µl of 800 µM dATP, dCTP, dGTP, 4SedTTP (final concentration of 80 µM for each), 2 µl RNase Inhibitor (80U final, NEB M0307L) and 2 µl SuperScript III (400U final, ThermoFisher 18080044) were added to each reaction to a final volume of 20 µl. The RT reactions were incubated at 42°C for 40 min, followed by incubation at 85°C for 15 min. The 100 nt long unmodified and m^6^A modified in vitro transcripts were treated in parallel with the experimental samples (Table 1). Negative controls included fragmented, affinity captured PAN RNA (at T0-T4 time point of infection) that was ligated to AR-17 oligonucleotide and directed to control RT reactions.

### Deep-sequencing data deconvolution

The libraries were prepared by the RNA ligation method (46) and paired-end sequenced on Illumina HiSeq 2500 platform (Psomagen USA), resulting in ∼1 million reads per sample. The trimmed (47) high-quality reads were mapped against PAN sequence using Bowtie2 (48). Mapped reads for each library pair (+4SedTTP, -4SedTTP) were scaled to the data set with the lowest number of mapped reads. Two metrics were used to analyze 4SedTTP RT sequencing data: 1) m^6^A_RATIO_, which describes the proportion of reads supporting RT stops at a position out of all the reads overlapping it, as per formula (1).

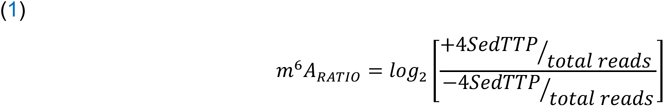

where +4SedTTP corresponds to RT reactions including 4SedTTP, and -4SedTTP corresponds to RT reactions including dTTP; 2) m^6^A fold change (m^6^A_FC_), which is the log_2_ transformed m^6^A ratio in the treated samples (+4SedTTP) divided by the m^6^A ratio in the non-treated samples (-4SedTTP) (23), as per formula (2). The reported m^6^A_FC_ value corresponds to the RT stop 1 nt upstream of the m^6^A position. (2)

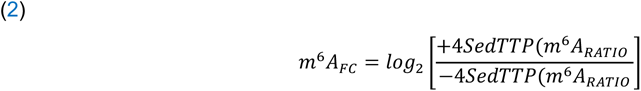

### Direct RNA sequencing analysis

The 500 ng of total RNA extracted from BCBL-1 cells at the KSHV latent (T0), and late lytic infection (T3) were subjected to ribosomal RNA depletion using the RiboCop rRNA Depletion Kit (Lexogen 037). The direct RNA sequencing library preparation was performed according to the manufacturer’s directions (ONT SQK-RNA002). Base-calling was performed using Albacore 1.2.1 [-f FLO-MIN106 -k SQK-RNA001 -r -n 0 -o fastq, fast5]. Reads present in the “pass” folder were used in the subsequent analysis. EpiNano software was utilized to determine the position of m^6^A (21). The SquiggleKit was used to view current intensities (22). Samtools and bcftools (49) were used for evaluating the number of insertions/deletions (INDELs), mismatches, and base quality scores.

### PAN RNA crosslinking (UV and FA) followed by antisense purification with mass spectrometry (RAP-MS)

2 x 10^7^ BCBL-1 cells per time point were crosslinked with 2% formaldehyde (VWR M134) in prewarmed 1X phosphate buffered saline (PBS) and incubated for 10 min at 37 °C (45). 2 x 10^7^ BCBL-1 cells per time point were crosslinked with UV light at 254 nm wavelength with 8000 x 100 µJ/cm^2^ in 1X PBS (50). These reactions were quenched with glycine to a final concentration of 500 mM and washed thrice with 1X PBS followed by PAN affinity capture. 60 pmol of antisense probes were added to 300 µl 1X hybridization buffer (20 mM Tris-HCl pH 7.5, 7 mM EDTA, 3 mM EGTA, 150 mM LiCl, 1% NP-40, 0.1% sodium deoxycholate, 3 M guanidine thiocyanate, 2.5 mM tris(2-carboxyethyl) phosphine (TCEP)), 5 µl protease inhibitor cocktail set (EDM Millipore 539134), and 10 µl murine RNase inhibitor (200U, NEB M0314L) and incubated for 2 h at 37 °C, after which they were washed six times with 1X PBS at 45°C. The elution of PAN-protein complexes was performed using 0.5 μl Benzonase nuclease (125U final, EDM Millipore 71206) in 1 ml of 0.35 M NaCl for 2 h at 37 °C (45, 50, 51).

### MS data processing and normalization

Captured proteins were analyzed using the Ultra-High-Performance Liquid Chromatography directed to Q Exactive Hybrid Quadrupole-Orbitrap Mass Spectrometry (YPED Proteomics at Yale School of Medicine). The Orbitrap raw files were analyzed using MASCOT (ver. 2.6) and the UniProt canonical human protein database supplemented with common contaminants. Samples were searched, allowing for a fragment ion mass tolerance of 10 ppm and cysteine carbamidomethylation (static) and methionine oxidation (variable). A 1% false discovery rate for both peptides and proteins was applied. Up to two missed cleavages per peptide were allowed, and at least four peptides were required for protein identification and quantitation with a fragment mass tolerance of 0.02 Da and a monoisotopic weight. Proteins that were specifically captured as interacting with PAN by both formaldehyde (FA) and UV crosslinking and that were not identified in non-crosslinked samples were directed for further analysis. Data normalization was performed using the R package NOMAD (Normalization of Mass Spectrometry Data (52) followed by PCA (Principal Component Analysis) to provide a dimensionality-reduction solution to identify the proteins that have a high confidence (p>0.05) for FA and UV crosslinking methods.

### Sodium dodecyl sulfate-polyacrylamide gel electrophoresis (SDS-PAGE) and Western blotting

2 x 10^7^ BCBL-1 cells were washed with 1X PBS and resuspended in Nuclear Lysis Buffer I (10 mM HEPES pH 7.4, 20 mM KCl, 1.5 mM MgCl_2_, 0.5 mM EDTA, 1 mM TCEP, 0.5 mM phenylmethylsulphonyl fluoride (PSMF) and 0.1% NP-40) supplemented with 10 µl of murine RNase inhibitor (200U final), and 5 µl protease inhibitor cocktail set (EMD Millipore 539134) in a total volume of 1 ml. Cells were sonicated for 180 sec at 4 °C, 15 peak power, 10 duty factor, 200 cycles at 1.5 average power with the wavelength adaptor no. 500534. DNase I (NEB M0303) treatment was performed according to the manufacturer’s protocol, followed by 30 min 16,000 x g centrifugation at 4 °C. The whole-cell protein lysates and the proteins crosslinked to PAN were analyzed by 10% SDS-PAGE. Proteins were transferred to polyvinylidene fluoride (PDVF) membrane using a semi-dry blotting system (ThermoFisher IB24002). The membranes were washed with 1X PBS with 0.1% Tween 20 (PBST) and blocked with 4% (w/v) non-fat milk in 1X PBST for 1 h at room temperature. The membranes were incubated with a specific primary antibody (Table 2) for 1 h at room temperature, washed thrice with 1X PBST, and incubated with the horseradish peroxidase (HRP)-conjugated secondary antibody (Table 2) for 45 min at room temperature. Blots were developed using the enhanced chemiluminescence substrate (BioRad 1705061).

**Table 2.** Primary and secondary antibodies used in this study.

| Primary antibody | Dilution for Western blot | Dilution for immunofluorescence analysis | Secondary antibody |
| --- | --- | --- | --- |
| RBM15 Polyclonal (ThermoFisher, PA5-22067) | 1:1000 | 1:50 | goat anti-rabbit immunoglobulin conjugated to horseradish peroxidase (Enzo, ADI-SAB-300-J) |
| METTL3 (ThermoFisher, MA5-34248) | N/A | 1:50 |  |
| METTL3 (ThermoFisher, MA5-27527) | 1:1000 | N/A | goat anti-rabbit immunoglobulin conjugated to horseradish peroxidase (Enzo, ADI-SAB-300-J) |
| FTO (ThermoFisher, PA5-51074) | 1:1000 | 1:200 | goat anti-rabbit immunoglobulin conjugated to horseradish peroxidase (Enzo, ADI-SAB-300-J) |
| hnRNP-C (ThermoFisher, PA5-22280) | 1:1000 | 1:200 | goat anti-rabbit immunoglobulin conjugated to horseradish peroxidase (Enzo, ADI-SAB-300-J) |
| YTHDF2 (Cell Signaling Technologies, 80014) | 1:1000 | 1:50 | goat anti-rabbit immunoglobulin conjugated to horseradish peroxidase (Enzo, ADI-SAB-300-J) |
| SND1 (Protein Technologies, 10760-1-AP) | 1:1000 | 1:200 | goat anti-rabbit immunoglobulin conjugated to horseradish peroxidase (Enzo, ADI-SAB-300-J) |
| Tubulin (ABCAM, Ab185224) | 1:1000 | N/A | goat anti-rabbit immunoglobulin conjugated to horseradish peroxidase (Enzo, ADI-SAB-300-J) |
| Actin - Mouse | 1:1000 | N/A | goat anti-mouse immunoglobulin conjugated to horseradish peroxidase (Abcam, ab97023) |

### siRNA knockdowns

1 × 10^7^ BCBL-1 cells were seeded in 30 ml fresh RPMI-1640 medium supplemented with 1% L-glutamine without antibiotics and without FBS to synchronize cells at G_0_ before transfection. For transfection, 80 pmol target-specific siRNAs (Santa Cruz Biotechnology, siMETTL3 sc-92172, siFTO sc-75002, siRBM15 sc-76365, and control siRNA-A sc-37007) were combined with 1 ml OPTI-MEM (ThermoFisher 31985062) and incubated for 5 min at room temperature. Lipofectamine RNAiMAX (ThermoFisher 13778) was combined with 1 ml OPTI-MEM and incubated for 5 min at room temperature. Both siRNAs/OPTI-MEM and Lipofectamine/OPTI-MEM solutions were combined, incubated for 5 min at room temperature, and added to the cells followed by 5 h incubation at 37 °C and 5% CO_2_. Following the incubation, the second siRNA-directed knockdown procedure was performed as described above. 24 h after transfection, 10% FBS was added to induce the cell cycle re-entry, followed by NaB treatment to the final concentration of 0.3 mM. The efficiency of siRNA knockdowns was verified by RT qPCR and Western blotting.

### Selenium-modified deoxythymidine triphosphates (SedTTP)-RT and Ligation Assisted PCR (SLAP)

2 x 10^7^ BCBL-1 cells per time point were used for total RNA extraction. The 5’ end phosphorylation was performed by treating 10 μg total RNA with 0.5 μl of RNase inhibitor (20U final, NEB M0307L), 1 μl of 10x T4 PNK reaction buffer, 0.1 mM ATP, 1 μl of T4 Polynucleotide kinase (10U final) in 10 μl total volume for 30 min at 37 °C. To ligate the RNA-5 oligonucleotide (Table 1), 1 μl of 10x T4 RNA Ligase Reaction Buffer, 100 pmol RNA-5 oligo, 0.1 mM ATP, 1 µl of RNase inhibitor (10U final), 3 μl of 100% DMSO, and 1 μl of T4 RNA ligase I (10U final, NEB M0437M) were combined in 20 μl total volume and incubated for 16 h at 16 °C. 13.5 μl of the ligation reaction was combined with 10x annealing buffer (250 mM Tris-HCl, pH 7.4, 480 mM KCl) and 0.5 pmol target-specific RT primer in 15.5 μl, and the reaction was incubated for 2 min at room temperature. 2 µl of 4SedTTP reaction buffer (500 mM Tris-HCl pH 8.0, 500 mM KCl, 50 mM MgCl_2_, 100 mM DTT), 2 µl of 800 µM dATP, dCTP, dGTP, 4SedTTP (final concentration of 80 µM for each), 2 µl RNase Inhibitor (80U final, NEB M0307L) and 1 µl AMV RT (12U final, ThermoFisher 18080044) were added to each ligation reaction to the final volume of 20 µl and incubated for 1 h at 42°C, followed by the addition of 1 µl RNase H (5U final, NEB M0297L) and incubation for 20 min at 37°C. To anneal the adapters, 1.5 pmol adapter/splint oligos mixture (1.5 μM each) was added to the RT mixture, followed by incubation for 3 min at 75°C. To ligate the adapter, 4 μl of T4 DNA ligase (40U final, NEB M0202L), 1X T4 DNA ligase reaction buffer, and 2 μl 100% DMSO were added to the above mixture in a total volume of 25 μl and incubated for 16 h at 16 °C. The reactions were purified using SILANE beads. PCR was performed with 0.2 µl Platinum Taq High Fidelity DNA Polymerase (2U final, ThermoFisher 11304011) in 25 μl total and the products were analyzed on 10% polyacrylamide gel electrophoresis (PAGE) and stained with 1X ethidium bromide solution overnight at 4 °C (53). Densitometric analysis of the PAGE gels was performed using ImageJ v1.52a (54) on inverted TIFF files.

### Selective 2-hydroxyl acylation analyzed by primer extension and mutational profiling (SHAPE-MaP) of PAN RNA

Each reaction included 1 × 10^7^ BCBL-1 cells washed with 1X PBS, collected by centrifugation at 300 × g for 5 min, and resuspended in 900 µl of RPMI-1640 medium. The SHAPE (+) reaction included the addition of 100 µl of 100 mM 1-methyl-7-nitroisatoic anhydride (1M7, DC Chemicals DC8649) resuspended in 100% DMSO to a final concentration of 10 mM to 900 µl of cells, followed by the incubation for 5 minu at 37°C. For the SHAPE (-) reaction, 100 µl of 100% DMSO was added to 900 µl of cells and incubated for 5 min at 37°C. The denatured control (dc) reaction included 1 µg/µl of total RNA extracted from BCBL-1 cells, that was combined with 5 µl 100% formamide and 1 μl 10X denaturing buffer (500 mM HEPES pH 8.0, 40 mM EDTA) in a total volume of 9 μl and incubated for 1 minute at 95 °C. 1 µl of 100 mM 1M7 was subsequently added to 9 µl of sample and incubated for 1 minute at 95 °C. SHAPE-MaP libraries were constructed by dividing the PAN RNA sequence into 3 overlapping zones, including nt 15-343 for zone 1, nt 311-707 for zone 2, and nt 694-1060 for zone 2 (Table 1) (20). Libraries were sequenced using MiSeq Nano V2 (Illumina) paired-end sequencing 2×250 bp with on average 100,000 reads per sample. SHAPE-MaP data were analyzed using the ShapeMapper v1.2 pipeline (21).

### Fluorescence in situ hybridization, co-staining, and image quantification

1 x 10^7^ BCBL1 cells were collected at the indicated time points post-induction, fixed with 4% paraformaldehyde (Electron Microscopy Sciences 15852-15s), permeabilized with 0.5 % Triton X, and washed thrice with 1X PBS. PAN specific oligonucleotide probes were labeled with a DIG-Oligonucleotide tailing kit (Sigma Roche 2^nd^ generation 03353583910) and mixed at the 1:1:1 ratio. Probes were denatured for 5 min at 95 °C before their addition to the hybridization solution (10% w/v dextran sulfate, Sigma D-6001), 10% v/v deionized formamide (VWR 97062-008), 2X nuclease-free saline sodium citrate), followed by the overnight incubation at 37 °C. After 1 h blocking, primary mouse anti-DIG (Sigma Roche 11333062910) was added at the dilution of 1: 200 in 1 % BSA and incubated for 1.5 h at room temperature. After three complete washes, the slides were incubated with a secondary Alexa Fluor 488-conjugated goat anti-mouse antibody (ThermoFisher A-11029) at a dilution of 1:500 in 1% BSA. Methylome detection including METTL3, RBM15, FTO, YTHDF2, HNRNPC, SND1 was performed by using specific primary anti-rabbit antibodies in 1% BSA (Table 2). After three complete washes, the secondary Alexa Fluor 647-conjugated goat anti-rabbit antibody (ThermoFisher A-21244) was added at a dilution of 1: 500 in 1% BSA. 1 µg/ml DAPI (ThermoFisher 62248) was used to stain nuclei for 30 min at 37 °C. The cells were mounted using VECTASHIELD® Antifade Mounting Medium H-1200 (Vector Laboratories) and observed under Nikon ECLIPSE Ti confocal microscope. Image J 1.52a was used to analyze FISH and IF images. To determine the raw fluorescence intensity in the stacked images, the embedded line tool and Plot Profile function were selected in Image J. The line traces quantitatively representing the raw fluorescence signals across all channels and data for the different stains were exported to Microsoft Excel, normalized, and plotted.

### Motif analysis

The identification of m^6^A consensus motifs within PAN RNA sequence and cellular lncRNAs sequences was performed with Multiple EM for Motif Elicitation (MEME, v5.1.1) using 8 nt windows surrounding the m^6^A site.

### RiboSNitch analysis

The analysis of PAN RNA structural changes invoked by the ablation of METTL3 and FTO was performed by using SNPfold (55) with set default parameters, followed by the variant analysis of global and local structural change by classSNitch (56).

### Data Visualization

Graphics were created using Microsoft Excel, Biorender (biorender.com), BEG Venn Diagram (bioinformatics.psb.ugent.be/webtools/Venn), Adobe Photoshop (ver 22.2), ImageJ (ver. 1.51), Geneious Prime 11.1.5 (https://www.geneious.com), Cytoscape ver. 3.8.2, and R studio ver. 3.6.2 (57).

## RESULTS

### PAN RNA is most extensively modified during the late lytic stages of KSHV infection

Most m^6^A mapping studies have been focusing on analyzing the global epitranscriptome of a specific organism or a virus with little insight into the position and frequency of modification on an individual transcript. The critical involvement of PAN RNA in the modulation of viral replication and cellular processes prompted us to examine whether this viral non-coding transcript undergoes m^6^A modifications, and if its epitranscriptomic status is dynamic. We designed a set of ten antisense oligonucleotide probes to affinity capture PAN RNA at the latent (T0), immediate early (T1 = 8 h post-induction, h pi), early (T2 = 24 h pi), and late lytic stages of replication (T3 = 48 and T4 = 72 h pi) (Supplementary Fig. 1a), which are defined by distinct gene expression programs in the KSHV infectivity cycle (5). The region of the KSHV genome from which PAN RNA is transcribed overlaps the K7/survivin locus (58). To differentiate between the epitranscriptomic imprints of PAN RNA and K7 mRNA, we designed an antisense oligonucleotide probe that targets the K7 transcript in a region upstream of the overlapping sequence (Supplementary Fig. 1b) (59). Additionally, the analysis included the antisense oligonucleotide probes specific to U1 small nuclear (sn) RNA to control for the affinity capture efficiency and specificity (51) (data not shown).

Previously it has been reported that the inclusion of 4-selenothymidine-5’-triphosphate during reverse transcription (4SedTTP RT) can identify the position of m^6^A on target RNA at single-nucleotide resolution (60). The presence of this deoxythymidine triphosphate (dTTP) derivative that carries a selenium atom replacing an oxygen at position 4 weakens its ability to base pair with m^6^A, creating a stop at the position immediately downstream of the modified site (+1) while maintaining normal A-T base pairing. We verified the accuracy of this approach by performing reverse transcription (RT) in the presence and absence of 4SedTTP on a 40 nt long transcript that carried a single m^6^A. The analysis confirmed the presence of a modification-induced truncation (Supplementary Fig. 2a). Next, we sought to determine the compatibility of the 4SedTTP RT method with next-generation sequencing analysis. We used two control transcripts, one carrying m^6^A marks at positions 13 and 82 and another unmodified transcript. The transcripts were directed to the 4SedTTP RT and control RT reactions, which instead of 4SedTTP, included thymine triphosphate (dTTP). To account for the m^6^A positional effect, we included random hexamer primers in all RT reactions. Out of 8,394 total reads obtained for the modified transcript in 4SedTTP RT reaction, 1,487 carried RT stops at the expected positions, resulting in the estimated m^6^A ratio of 5.6% (m^6^A_RATIO_, defined as the proportion of reads supporting RT stops at a position out of all the reads overlapping it). No reads indicating truncations were detected for control RT reactions performed on modified (dTTP-m^6^A), unmodified transcript (dTTP-A), and the 4SedTTP RT reaction performed on unmodified transcript (4SedTTP-A). The m^6^A fold change (m^6^A_FC_, defined as the log_2_ transformed m^6^A ratio in the treated samples (+4SedTTP) divided by the m^6^A ratio in the non-treated samples (-4SedTTP)) for two sites on modified transcript (4SedTTP-m^6^A), was estimated at 8.5 and 8.1, respectively (Supplementary Fig. 2b). The m^6^A_FC_ calculated for 4SedTTP-m^6^A/dTTP-m^6^A libraries showed no significant correlation between the results, while that same ratio calculated for 4SedTTP-A/dTTP-A libraries indicated significant correlation (*ρ* = 0.99%, pairwise *p*-value <10^-13^). This indicated that RT stops, other than the ones induced by modification, were random and nonspecific.

After verifying the accuracy of the 4SedTTP RT method, we used it to map m^6^A on PAN RNA captured at the latent (T0) and lytic stages of KSHV replication (T1 – T4). We identified a total of five peaks, that mapped to m^6^A residues at nt positions 18 and 203, 672, 1041, and 1048 (Fig. 1). These results were reproducible across two distinct biological replicates. The peak representing the modification at nt 18 showed up first during the latency and became pronounced at the late lytic stage of KSHV infection (T4). The peak corresponding to m^6^A at nt 203 was weak at the immediate early lytic stage (T1) and became pronounced at both late lytic stages of KSHV infection (T3-T4). The peak corresponding to m^6^A at nt 672 was detectable during the latency and became pronounced at the immediate early (T1) and late lytic stages of KSHV infection (T3-T4), while at the early lytic stage (T2), it was not detectable. The peaks at nt 1041 and 1048 were both the most distinct at the late lytic stages (T3 - T4) (Fig. 1). These data demonstrated that the PAN m^6^A landscape is dynamic and changes during the KSHV infectivity cycle. They also indicated that PAN RNA expressed at the late lytic stages of infection is the most extensively modified, as it can carry up to five m^6^A sites.

**Fig. 1.**
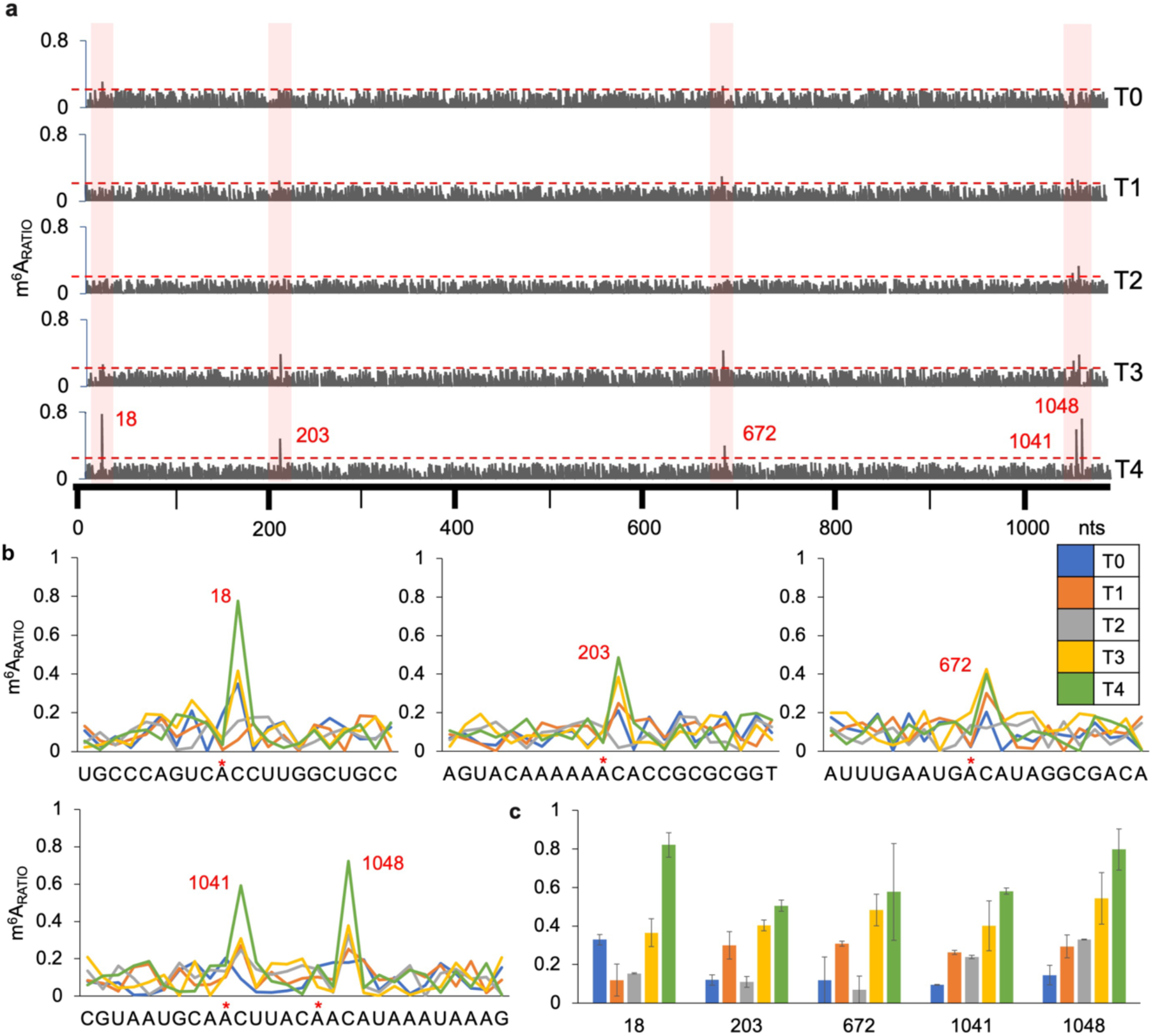
The m^6^A landscape of PAN RNA expressed at latent and lytic stages of KSHV replication. **a**, The graphs represent m^6^A_RATIO_ (Y-axis) against PAN RNA sequence (X-axis). The peaks correspond to the expected RT stop 1 nt upstream of m^6^A. The threshold line (red) for calling m^6^A peaks is set at 0.2 and it was calculated by dividing the average m^6^A_RATIO_ values calculated for all PAN nucleotides divided by the total number of nucleotides. **b**, The charts represent the signal-to-noise ratio for sequences overlapping m^6^A peaks within specified nucleotide window (X-axis). **c,** The average m^6^A_RATIO_ values for each modification at specified time points. Time points of KSHV infection are color-coded as follows: latency - T0 in blue; lytic stages: immediate early – T1, 8 h post infection (h pi) in orange; early - T2, 24 h pi in gray; late - T3, 48, h pi in yellow; late - T4, 72 h pi in green.

### Direct RNA sequencing (DRS) confirms m^6^A modifications on PAN RNA

The DRS has been adopted to examine the m^6^A status of full-length RNAs (61–63). It has been shown that the base calling of reads corresponding to m^6^A results in a higher mismatch frequency (MF) as compared to the unmodified reads (64). Also, base quality (BQ), insertion and deletion (INDEL) frequency, and current intensity can be affected by the presence of m^6^A (64). We sought to determine these values for PAN RNA expressed during KSHV latency (T0) and at the late lytic stage of replication (T3), during which the transcript carried the largest number of modified residues (Fig. 1) and compared them to two control transcripts. We found that the mismatch frequency in the 100% modified transcript was significantly higher than that of the unmodified transcript (*p*-value for MF = 1.868×10^-135^, t-test, *p*-value < 0.05) (Fig. 2a). A similar correlation was noted for PAN expressed during KSHV lytic replication versus latency (*p*-value for MF = 1.586×10^-47^, t-test, *p*-value < 0.05). The analysis of mismatch directionality for m^6^A in 0% and 100% modified control transcripts, and PAN expressed during latency (T0) and late lytic stage (T3) of KSHV replication indicated that the modification did not invoke biases towards any specific type of mismatch (Supplementary Fig. 3a). Also, we did not observe that the INDEL frequency, base quality scores, or current intensity were significantly different among analyzed samples (Supplemental Fig. 3b-d).

**Fig. 2.**
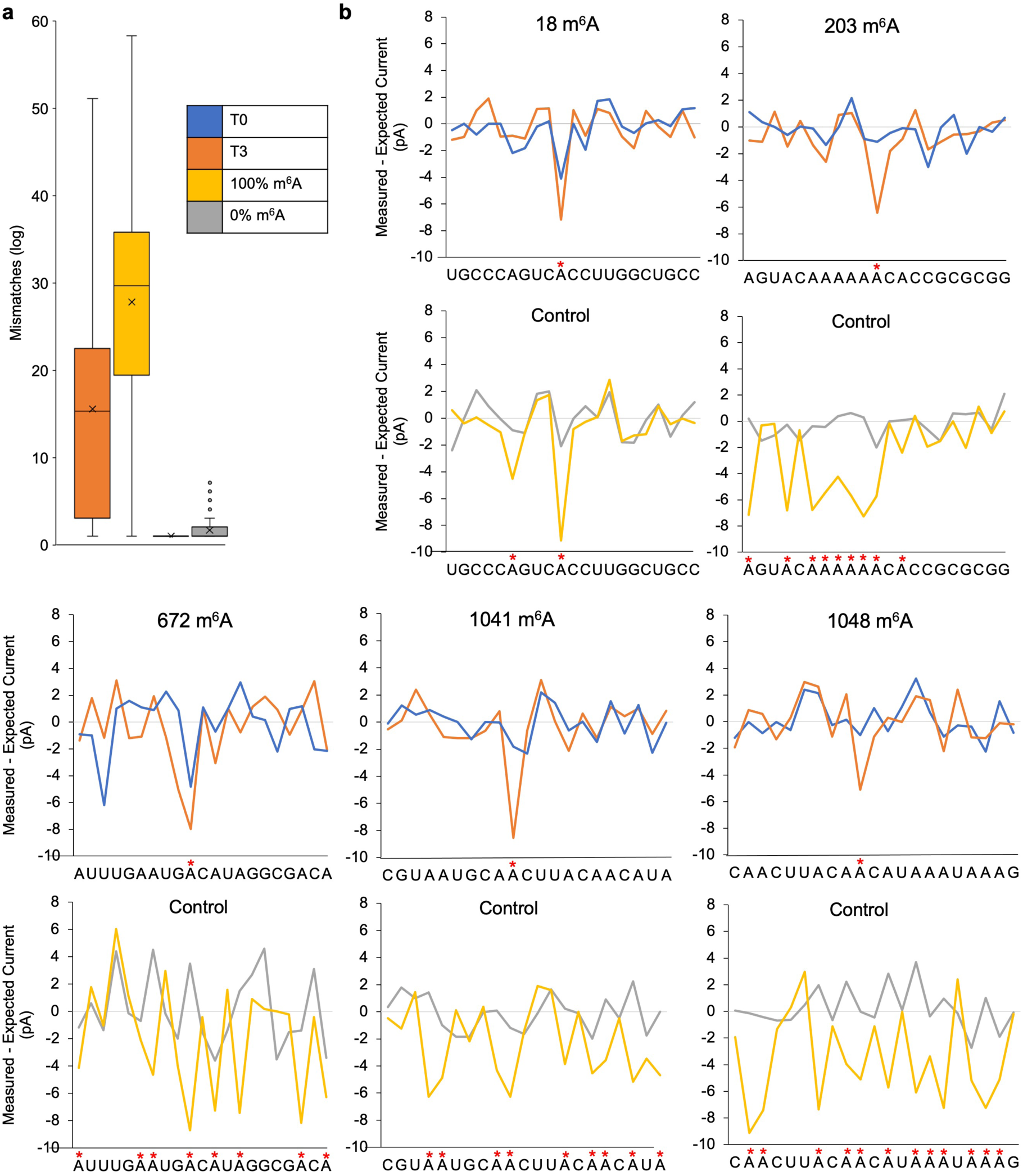
The results of DRS data analysis for PAN RNA. **a**, Mismatch frequency (MF), **b**, The presence of m^6^A on PAN RNA expressed during KSHV latency (T0, blue), and late lytic stage (T3, orange) induces the distortion in electric current. Similar current distortions are noted for all m^6^A on 100% modified PAN transcript (yellow) versus unmodified transcript (gray).

Previous studies have shown that DRS raw current intensity signals can be used to identify the current changes invoked by the presence of m^6^A with single-nucleotide resolution (61, 65). In particular, the EpiNano, nanopore base-calling software, trained with m^6^A modified and unmodified synthetic sequences has been shown to recognize m^6^A with ∼90% accuracy in vitro, while in vivo modifications have been identified with ∼87% accuracy and an m^6^A recovery percentage of around 32% (defined as the number of positive methylation signals divided by the total number of adenosines) (64). We verified these findings by analyzing the current intensity signals obtained for 100% modified and unmodified PAN transcripts. Out of 259 m^6^A sites present in the modified transcript, we detected 183 current distortions, estimating the recovery percentage at 64% (Supplementary Fig. 3e). No current alterations were noted for unmodified transcript. We applied this approach to validate whether the presence of m^6^A on PAN expressed during the latent (T0) and lytic (T3) KSHV replication induces current distortions at modified sites. We mapped a total of ∼8000 reads corresponding to PAN RNA for each time point and assessed the recovery percentage for each m^6^A site. For PAN RNA expressed during latency, we observed the highest recovery percentages at sites 18 and 672 (18.4% and 11.2%, respectively), while the other sites had lower recovery rates (203 – 1.0%, 1041 – 3.3%, 1048 – 4.2%, respectively). For PAN RNA expressed during the late lytic KSHV replication, all PAN m^6^A sites had high recovery percentages (sites 18 – 35.3%, 203 – 29.5%, 672 – 15.3%, 1041 – 40.9%, and 1048 – 45.3%).

To validate the precision of the m^6^A-calling by EpiNano and visualize the results, we used the MotifSeq (SquiggleKit pipeline). The MotifSeq takes a primary query sequence as input (.fasta file), converts it to a normalized signal trace, and performs a signal level local alignment using a dynamic programming algorithm (66). The normalization of signal traces occurs by subtracting the expected current values from the observed currents gathered by MotifSeq. This approach is particularly useful for detecting m^6^A, as signal traces for modified residues have been shown to hold a greater disparity between expected and observed currents (62). We compared the current values obtained for modified control transcript, the latent, and lytic PAN RNA samples with the unmodified control sample. Two m^6^A at nt positions 18 and 672 on PAN RNA expressed during the KSHV latency triggered current alternations (Fig. 2b). For PAN RNA expressed during the late lytic replication (T3), all m^6^A sites were manifested with current changes. A similar pattern of distortion was seen differentiating the modified and unmodified transcripts.

### The modification frequency at identified m^6^A on PAN RNA reaches the highest level during the late lytic stages of KSHV infection

The modification abundance at a specific site is a variable function, that can be instrumental for its functional importance in the context of specific transcripts. Unfortunately, high throughput epitranscriptomic mapping approaches, which involve multiple chemical handling steps, often result in higher than desired background, that obscures that relevant measurement. Also, the quantification of m^6^A stoichiometry based on DRS readouts has limitations as detection of modifications is partly dependent on sequence context and stoichiometry prediction is heavily affected by choice of re-squiggling algorithms (67). To gain accurate insight into KSHV replication state-dependent variations in PAN m^6^A at each identified site, we developed a new quantitative method, termed <u>S</u>elenium-modified deoxythymidine triphosphate Reverse Transcription and <u>L</u>igation <u>A</u>ssisted <u>P</u>CR analysis (SLAP). In this method, we used the SedTTP nucleotide in an RT reaction to induce stops 1 nt upstream of m^6^A. The resulting truncated m^6^A-specific products are ligated to adapter sequence present in the full-length unmodified products to introduce a common forward primer binding site. A set of PCR primers, including forward primer being complementary to the adapter and reverse primer being complementary to the same sequence on both modified and unmodified product, is used to simultaneously amplify both products, yielding the quantitative information for individual sites of modification. We first evaluated the accuracy and sensitivity of the SLAP method by setting up a calibration curve on samples that included a total of 30 fmoles of in vitro synthesized m^6^A modified and unmodified PAN transcripts that were combined at specific percentages and spiked with 1 µg of total RNA extracts from BCBL-1 cells (Fig. 3a). For samples that carried a range of modification from 2.5 - 100%, linear regression reached R^2^ = 0.988 (Fig. 3b). We estimated from the calibration curve that SLAP allows for quantification of the modification fraction at as low as 2.5 percent level. We noticed the presence of a very weak m^6^A-specific product in the positive reaction with only the unmodified transcript (+0%, Fig. 3a), which was regarded as the background, and subsequently a 2-fold threshold above that background was used to estimate the modification frequency. Minor background products were also noted for negative (-) control reactions but only for some m^6^A sites, i.e., at nt 18, suggesting the occurrence of nonspecific-to-m^6^A truncations in some RT reactions. Interestingly, the sample consisting of 100% modified transcript resulted in an estimation of m^6^A frequency at 92 percent level, which suggests a slight underestimation of the actual modification fraction (Fig. 3b).

**Fig. 3.**
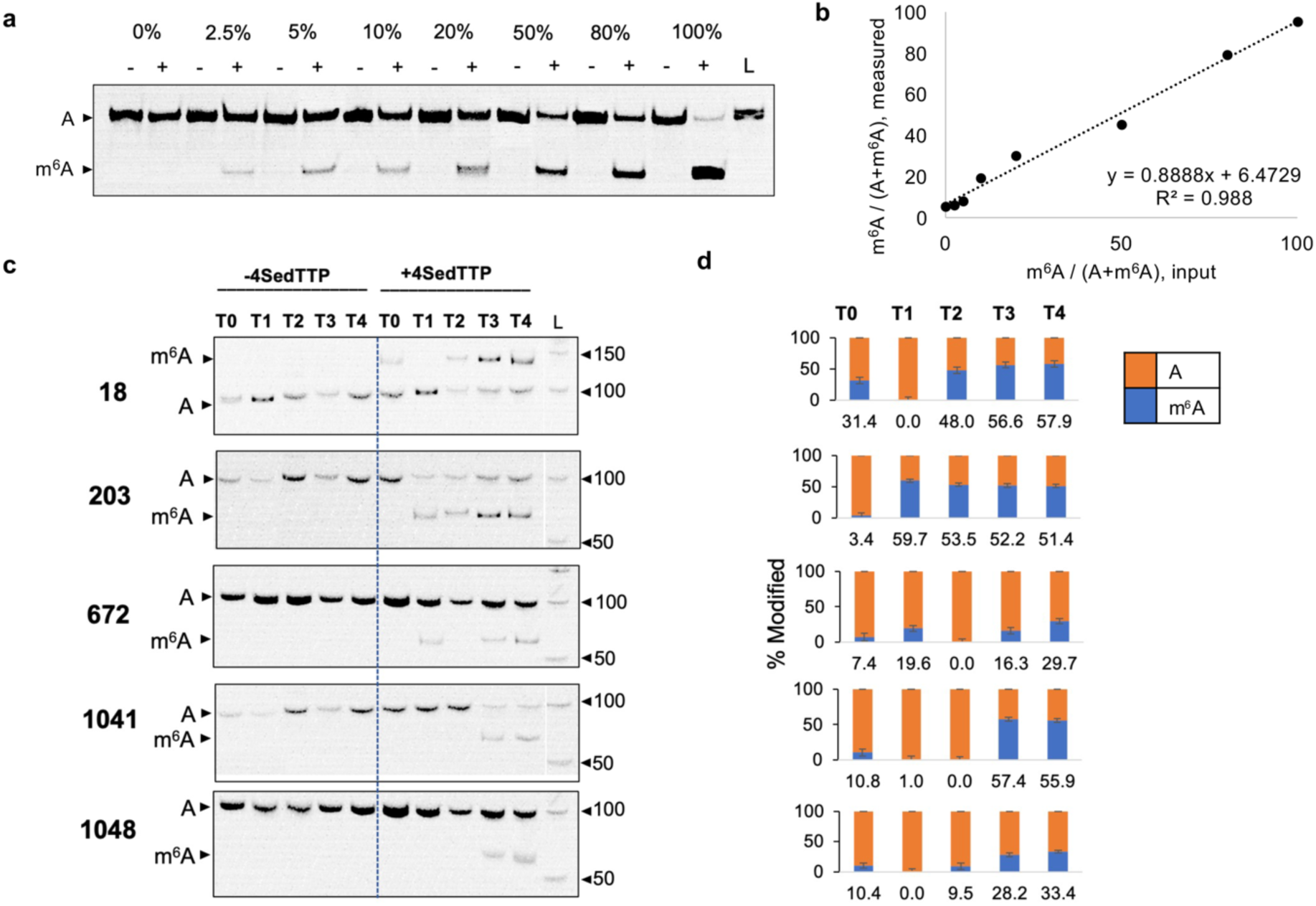
<u>S</u>elenium-modified deoxythymidine triphosphates Reverse Transcription and <u>L</u>igation <u>A</u>ssisted <u>P</u>CR analysis (SLAP) quantifies the modification frequency on PAN RNA. **a**, Native PAGE shows PCR products derived from the modified and unmodified RNA standards, combined at the indicated percentages and subjected to SLAP analysis. The negative (-) and positive (+) reactions are indicated on top, together with the percentage of modified RNA. **b**, The graph represents the SLAP standard calibration curve created based on quantification of results from panel A. The X-axis represents m^6^A fractions tested, while the Y-axis represents corresponding densitometric measurements (*n*=3) **c,** Native PAGE for representative replicate (n=3) of SLAP analysis performed for nt 18, 203, 672, 1041, and 1048 on PAN RNA. The time points of infection are indicated on top (T0 – T4). The position of products specific for modified and unmodified sites is labeled. L stands for 50 bp DNA ladder. **d,** The column graphs represent the average modification frequency on PAN at each site and time point of infection (*n* = 3). Standard deviations for frequency measurements are indicated.

Next, we applied the SLAP analysis to quantify the frequency of m^6^A at each identified site on PAN RNA (Fig. 3c). Due to the proximity of m^6^A at nt 18 to the 5’ terminus, analysis of that site required the use of extended adapter with the forward primer binding site that would allow differentiation of modified and unmodified products. As a result, the m^6^A-specific product corresponding to that site is longer (127 nts) than the product corresponding to unmodified residue (100 nt). Overall frequency of modification at all sites was relatively low for PAN RNA expressed during latent and early lytic stages of KSHV replication reaching the highest levels during late lytic stages (T3 – T4). In particular, the m^6^A at position 18 was estimated at 31.4±3.5% during latency, decreasing at immediate early stages to 0±1.7%, followed by an increase through lytic and late lytic phases to 48.0±2.9%, 56.6±2.7%, and 57.9±2.6%, respectively. The modification level at nt 203 was estimated at 3.4±4.8% during latency, 59.7±2.5% during immediate early, and 53.5±4.0% at early stages of viral replication, followed by an increase to 52.2±5.2%, and 51.4±2.8% during the late lytic stages (T3 – T4). Position 672 was modified at 7.4±5.0% during latency, 19.6±3.7% at immediate early, 0±5.1% at early lytic stages, followed by an increase to 16.3±4.2% and 29.7±3.2% during the late lytic stages of KSHV replication. Nucleotide 1041 was modified at 10.8±5.2% during latency, 1.0±1.6% during immediate early, 0±1.3% during early lytic stages, and showed the highest frequency during the late lytic stages, reaching a frequency of 57.4±5.3%, and 55.9±4.9%, respectively. The modification frequency at nt 1048 was detectable at 10.4±7.5% level during latency, reaching a frequency of 0±0.7% and 9.5±1.6% for immediate early and early lytic stages. However, the frequency of m^6^A at that site, again increased during the late lytic stages of KSHV replication, 28.2±9.8%, and 33.4±3.7%, respectively (T3 – T4).

The m^6^A modifications in human transcriptome have been primarily localized within two canonical motifs, Gm^6^AC (∼70%) and Am^6^AC (∼30%) (68, 69), although another study identifies GGm^6^AGG motif as present at the 41% frequency (70). Recent study performed on KSHV transcriptome has reported that approximately 60% of all m^6^A is contained within a GGm^6^AC[G/U] motif (37). We performed a motif search to investigate the m^6^A consensus sequence specific to PAN RNA. The overall PAN m^6^A consensus sequence was designed as A/G/Cm^6^ACA/U/C, showing higher sequence variability compared to the previously identified consensus (Supplementary Fig. 4). In particular, the m^6^A at position 18 was located within a Cm^6^AC motif, three modifications at nt positions 203, 1041, 1048 were located at an Am^6^AC motif, while the m^6^A at nt 672 was associated with a Gm^6^AC motif. We also analyzed the m^6^A consensus for cellular lncRNAs that have been previously reported as m^6^A modified. Here, we found that the majority of m^6^A modifications were located within an A/Gm^6^AC consensus sequence (Supplementary Fig. 4).

### PAN RNA associates with m^6^A writers, readers, and erasers

The RNA m^6^A status is dynamically coordinated by the catalytic activity of writer and eraser proteins, while the phenotypic effect of m^6^A is facilitated by various groups of reader proteins (71–74). To identify the methylome that is involved in the regulation of PAN m^6^A status, we performed crosslinking of PAN RNA-protein complexes during KSHV latent (T0) and lytic (T3) replication, followed by affinity capture, and mass spectrometry analysis (RAP MS) (75). RAP MS relies on 254 nm UV light crosslinking to create covalent bonds between RNA directly interacting with proteins (76, 77). The biggest caveat of UV crosslinking is its low efficiency, which has been estimated at ∼5% for any given ribonucleoprotein complex (78). Thus, we also performed formaldehyde (FA) crosslinking, which occurs through the formation of methylol derivatives between amino groups of cytosines, guanine, and adenines or the imino groups of thymines with amino acids such as lysine, arginine, tryptophan, and histidine (79). Both analyses were performed in three independent experimental replicates to account for methodological bias. We assessed the relative abundance of PAN RNA by RT qPCR and showed that the target was efficiently captured at both time points (over 100-fold enrichment for both T0 and T3, Supplementary Fig. 5a). The proteins that were captured in crosslinked samples and were not present in the non-crosslinked samples were compared statistically between methods and the replicate samples. This categorical analysis of our PAN-associating proteins and their mass spectrometry MASCOT scores showed that the number and type of proteins bore a strong correlation between the two crosslinking methods (FA and UV, R^2^ = 0.9894) and among distinct biological replicates (R^2^ = 0.9827 for replicates I and II, R^2^ = 0.9954 for II and III, R^2^ = 0.9901 for I and III, Fig. 4a). We also performed principal component analysis (PCA) on the data obtained from RAP MS with UV and FA crosslinking and found that the number and type of PAN-associating proteins shared 99% overlap between both crosslinking methods, and among the three distinct biological replicates (Fig. 4b).

**Fig. 4.**
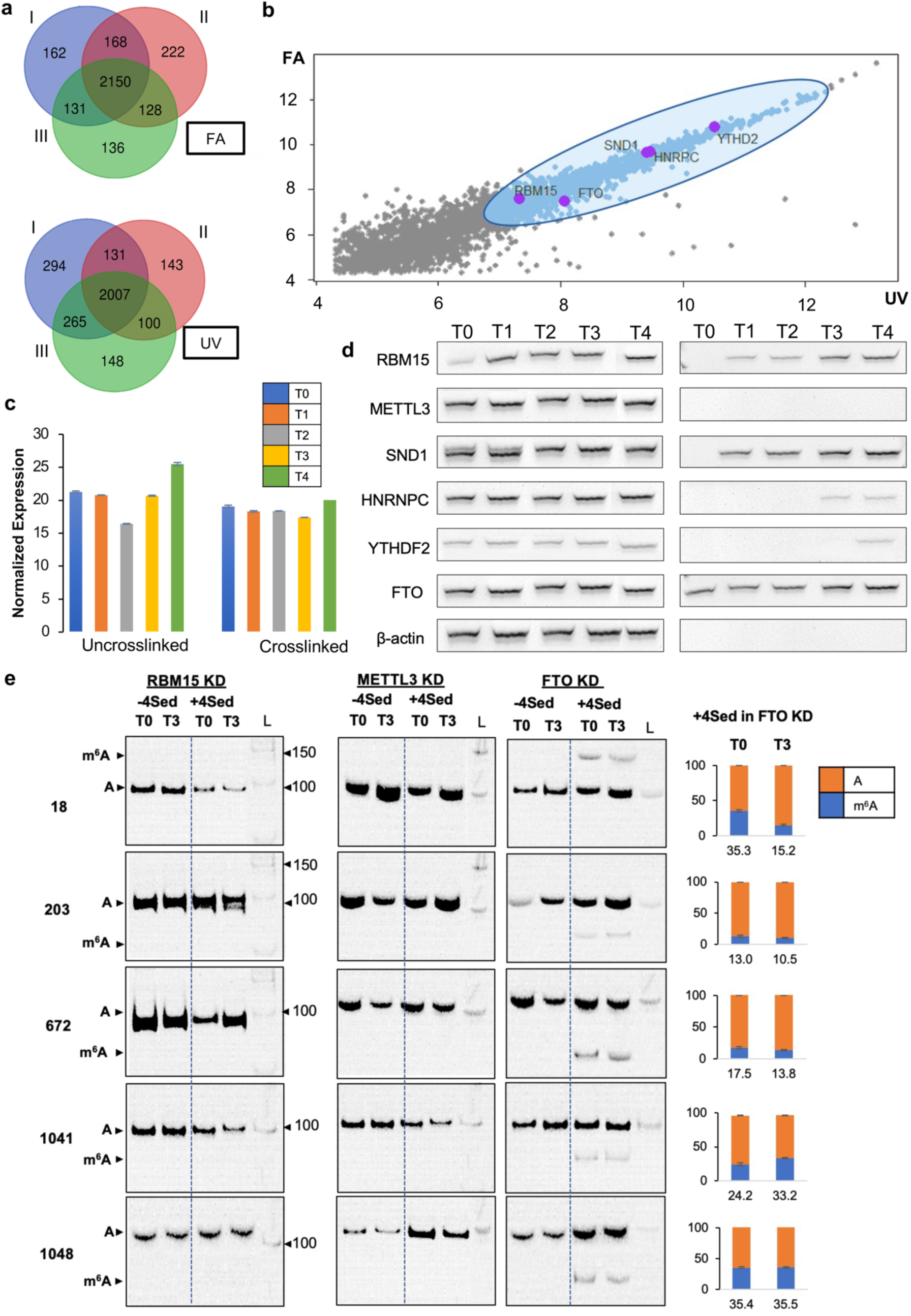
The RAP MS analysis for PAN RNA m^6^A methylome and effect on PAN m^6^A methylation frequencies. **a**, Venn diagrams representing the overlap between results generated from formaldehyde (FA) and UV crosslinked PAN-protein complexes. The three distinct biological replicates are indicated with colors (I-III). Normalized expression of PAN RNA assessed by RT qPCR in uncrosslinked total RNA samples extracted during KSHV latent (T0) and lytic (T3) stages of replication, and after the PAN RNA-protein crosslinking and affinity capture. **b**, Principal component analysis (PCA) of proteins identified in RAP MS (UV, X-axis) and FA experiments (Y-axis). Proteins involved in regulation of the PAN m^6^A status are marked with purple dots and they include the writer complex recruiter RBM15, readers HNRNPC, YTHDF2, SND1, and the FTO eraser. **c**, Normalized expression of PAN RNA in affinity captured fractions before (left) and after (right) crosslinking. **d**, Results of Western blot analyses performed on whole cells BCBL-1 lysates (left) and captured PAN RNA-protein complexes (right) during latent (T0) and lytic (T1-T4) stages of KSHV replication. β-actin was used as a control. **e**, The representative native PAGE indicating frequency of modification at m^6^A sites in BCBL-1 RMB15, METTL3, and FTO knockdown (KD) cells. L represents 50 bp DNA ladder. The analyzed time points of infection (T0, T3), negative (-4Sed) and positive (+4Sed) reactions are indicated on top. The position of products specific for modified and unmodified sites is indicated to the left. The column graphs represent the average modification frequency for each site at T0 and T3 time point of infection in specified knockdown cell lines. Standard deviations for frequency measurements are indicated (n = 2).

For PAN RNA expressed during the KSHV lytic replication (T3), we identified RBM15, a protein crucial for the recruitment of METTL3/14 writer complex (39), and demethylase FTO. We also identified three m^6^A reader proteins, i.e., HNRNPC, YTHDF2, and SND1. Importantly, none of these proteins were found to associate with PAN RNA expressed during the KSHV latency (T0). We established the lncRNA-<u>p</u>rotein interactions network (LPI) including the identified PAN methylome components and predicted interacting proteins derived from STRING, the functional protein association network database (80) (Supplementary Fig. 5b). The network identified 5 nodes (proteins) and 8 edges (interactions) associated with PAN m^6^A modification. Further, we classified RBM15 as a central hub for PAN m^6^A methylome, that was predicted to interact with the METTL3/14 writer complex, accessory proteins, i.e., RNA binding protein 15 (RBM15), Wilms’ tumor 1-associating protein (WTAP), and identified reader proteins, i.e., SND1, YTHDF2, HNRNPC. The FTO, PAN-associating eraser was predicted to bind METTL3, RBM15B, and two identified reader proteins, i.e., SND1 and YTHDF2. Overall, the LPI network was interconnected within each functional subset of PAN m^6^A methylome and between writers, erasers, and readers.

We further verified the associations of PAN RNA with these methylome components by performing a Western blot analysis. We first assessed the expression of individual methylome proteins in BCBL-1 whole-cell lysates and showed that their expression levels were not significantly affected by the progression of KSHV replication. After performing FA crosslinking and affinity capture of PAN RNA-protein complexes, we assessed the relative abundance of target RNA and showed that it was efficiently captured at all time points (Fig. 4c). We were able to confirm that all proteins identified by RAP MS analysis interact with PAN RNA. RBM15 and SND1 were shown to interact with PAN at all lytic stages of KSHV infection (T1-T4), but not during latency (T0). FTO was captured as interacting with PAN during all time points (T0-T4), while two reader proteins, HNRNPC and YTHDF2 were detected at the late lytic stage of KSHV infection (T3-T4, and T4, respectively) (Fig. 4c). METTL3 was not captured as directly interacting with PAN RNA, perhaps because, the METTL3/14 writer complex is known to interact with the target RNA via RBM15 association (81). We also assessed the normalized expression levels of mRNAs encoding the crosslinked proteins and showed that their expression levels varied marginally throughout the progression of KSHV infectivity cycle. Only YTHDF2 mRNA showed significantly increased expression during the lytic stages, however, that increase was not reflected in the expression of YTHDF2 protein (Supplementary Fig. 6).

### The knockdowns of methylome components affect PAN RNA expression levels and its m^6^A landscape

To assess the involvement of METTL3, RBM15 and FTO in the regulation of PAN m^6^A status, we performed the siRNA-directed knockdowns of individual proteins in BCBL-1 cells, followed by induction of KSHV lytic replication. The RT qPCR analysis indicated nearly complete expressional ablations of RBM15 (∼83%, Supplementary Fig. 7a), METTL3 (∼89%, Supplementary Fig. 7b), and FTO (∼92%, Supplementary Fig. 7c) at mRNA levels for each time point. Western blots further verified the efficiency of knockdowns for all three proteins (Supplementary Fig. 7d). It has been previously shown that an FTO knockdown in BCBL-1 cells induces expression of KSHV lytic genes in contrast to a METTL3 knockdown that abolishes lytic gene expression (36). We assessed the level of PAN RNA expression in the expressionally ablated cells and noted that for the FTO knockdown, the PAN levels initially increased (T1) after lytic induction, followed by sharp decline at the later stages of KSHV lytic replication (T2-T4) (Supplementary Fig. 7e). For METTL3 knockdown, there was a strong decrease in PAN expression at most time points of infection (T0-T3) with the exception of the late lytic stage (T4). In the case of the RBM15 knockdown, PAN RNA expression was decreased at all time points as compared to the levels observed for control BCBL-1 cells.

Next, we applied SLAP to validate the involvement of these methylome components in PAN modification. As anticipated, the knockdown of METTL3 and RBM15 completely abolished the installation of m^6^A on PAN RNA at all identified sites, and at both latent (T0) and late lytic (T3) stages of infection (Supplementary Fig. 8). Meanwhile, the FTO knockdown resulted in an m^6^A pattern similar to the one observed in wild-type BCBL-1 cells (Fig. 1). In particular, peaks of slightly lower m^6^A_RATIO_ were observed at nt 18 and 672 during latency (T0), and at all lytic stages of KSHV replication (T1-T4). We further verified these effects by performing SLAP analysis on PAN RNA expressed by each of the knockdown cell lines at latent (T0) and late lytic (T3) stages (Fig. 4e). Again, the m^6^A-specific products were not detected for METTL3 and RBM15 knockdowns at both analyzed stages of KSHV infection, and for FTO knockdowns, each site previously identified as modified resulted in generation of m^6^A-specific products. For site 18, the modification frequency reached 35.3±1.8% during latency (WT – 31.4±3.5%), and 15.2±1.3% during late lytic stage (WT – 56.6±2.7%). For site 203, the modification levels were 13.0±6.3% (WT – 3.4±4.8%) and 10.5±1.2% (WT – 52.2±5.2%) for latent and late lytic stages, respectively. For the residue at position 672, the modification levels were at 17.5±12.7% (WT – 7.4±5.0%) and 13.8±3.1% (WT – 16.3±4.2%), respectively. However, the modification abundance at sites 1041 and 1048 increased for both time points of KSHV infection and reached 24.2±2.7% (WT – 10.8±5.2%) and 33.2±1.3% (WT – 57.4±5.3%) for site 1041, and 35.4±4.2% (WT – 10.4±7.5%) and 35.5±1.0% (WT – 28.2±9.8%) for site 1048 (T0 and T3, respectively).

### The m^6^A modification affects PAN RNA secondary structure at local and global levels

We performed selective 2’-hydroxyl acylation analyzed by primer extension and mutational profiling analysis (SHAPE-MaP) of PAN RNA expressed in wild-type, METTL3 and FTO knockdown BCBL-1 cell lines to gain insight into the structural impact of m^6^A. We chose to probe the structure of PAN RNA expressed during the late lytic stage of infection (T3) as at that time point the transcript carried the largest number of modified residues (Fig. 1) at the highest modification frequency (Fig. 3). It has been shown that m^6^A methylation occurs at sites that are mostly un-paired (82, 83). Hence, we anticipated that the expressional ablation of METTL3 would trigger more double-stranded and constrained structure, while the knockdown of FTO would invoke more structurally flexible, i.e., un-paired regions in PAN RNA. Since probing of PAN RNA secondary structure in wild-type BCBL-1 cells (Supplementary Fig. 9) would be expected to yield the ensemble of reactivity values at m^6^A reflecting varying modification frequency and distribution, we compared the reactivity profiles (ΔSHAPE analysis) for PAN RNA expressed in METTL3 and FTO knockdown cells (Fig. 5a) (12, 84). As expected, we observed that the ablation of writer invoked lower reactivity values at m^6^A sites, while the knockdown of eraser induced strong structural signatures marked by increased exposure of m^6^A residues to the modifying reagent. The noted structural changes were of two type - local, defined as within 20 nt to the identified m^6^A (nt 25 - 32, 195 - 210, 703 - 709, and 1039 - 1050), and global, defined as outside the 20 nt range (nt 120 - 130, 151 - 158, 820 - 827, 900 - 950) (Fig. 5a). The most pronounced global changes invoked by the ablation of METTL3 were noted within the <u>e</u>xpression and <u>n</u>uclear retention <u>e</u>lement (ENE), which is adjacent to two m^6^A at nt 1041, and 1048 (85). The PAN ENE forms a hairpin containing a U-rich internal loop flanked by short helices, which is involved in the sequestration of poly(A) tail, leading to the formation of stabilizing triple helix (85, 86). Interestingly, in METTL3 knockdown cells, the ENE U-rich residues were split between two hairpins (Fig. 5b, c). Such extensive secondary structure rearangement was not noted for PAN RNA expressed in the FTO knockdown BCBL-1 cells, which was predicted to invoke mostly local structural rearagnements.

**Fig. 5.**
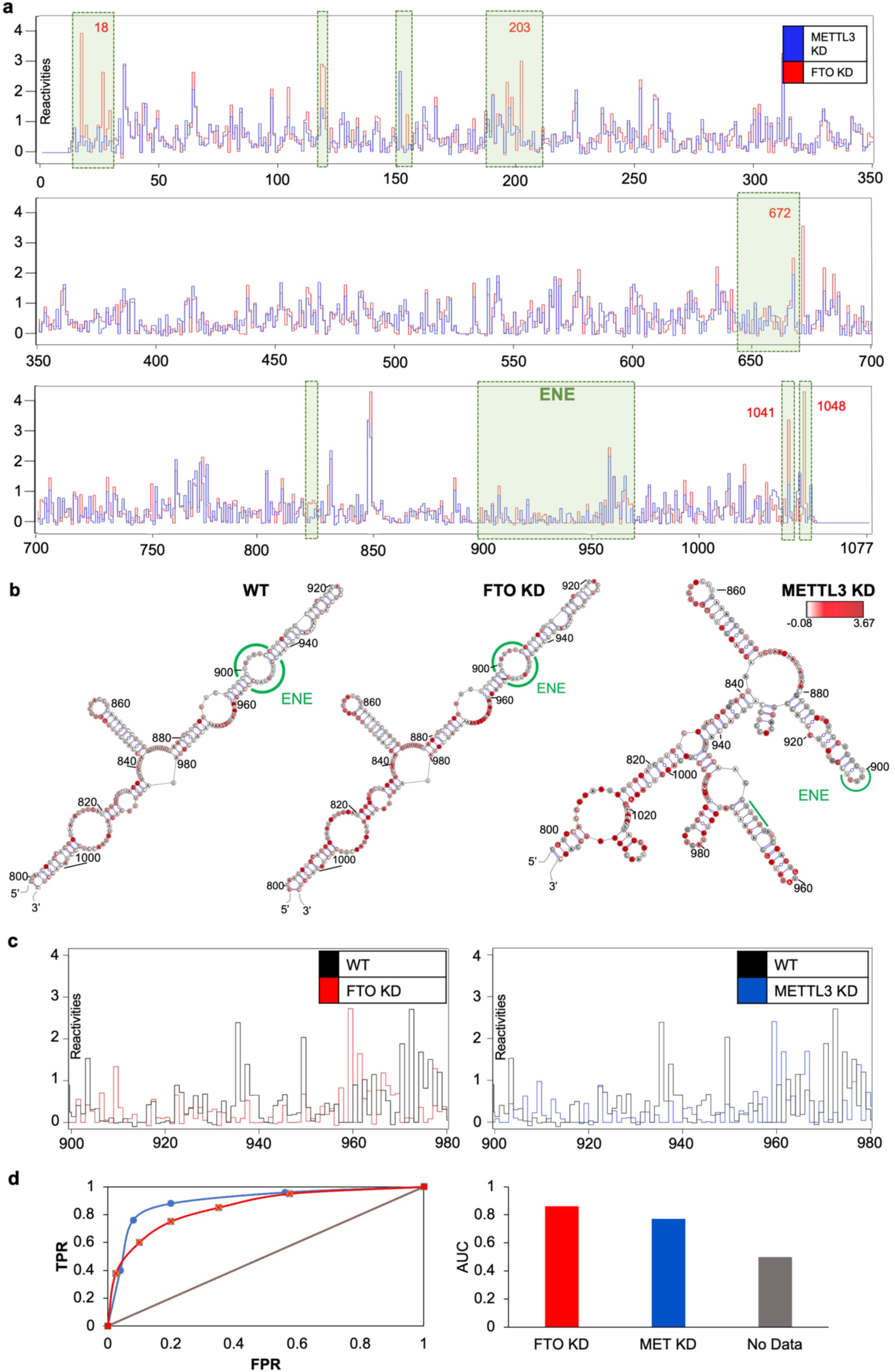
The structural analysis of PAN RNA in METTL3 and FTO knockdown BCBL-1. **a**, PAN RNA SHAPE-MaP reactivity profiles with blue trace corresponding to reactivity values obtained in METTL3 KD cells, red trace representing reactivity values from FTO KD cells. X-axis indicates nucleotide position on PAN RNA, Y-axis represents SHAPE-MaP reactivity values. Nucleotides 1-14 and 1056-1077 were not covered in the analysis due to primer annealing. The green boxes indicate the local and global structural changes invoked by the methylome ablations. The position of the <u>e</u>xpression and <u>n</u>uclear retention (ENE) motif is indicated. **b**, The secondary structure of PAN ENE motif (green) probed in wild-type, FTO and METTL3 knockdown (KD) cells. Nucleotides are colored by SHAPE reactivity as follows: grey residues display low reactivity (< 0.2), pink residues representing medium reactivity (0.2 – 0.8), and red values representing high reactivity (> 0.8). **c**, The comparison of PAN ENE reactivity values read out in wild-type and FTO KD BCBL-1 cells and wild-type and METTL3 KD BCBL-1 cells. **d**, ROC curves (left) generated based on SHAPE reactivity values and representing the m^6^A influence over PAN RNA secondary structure. The X-axis corresponds to False Positive Rate (FPR), the Y-axis represents True Positive Rate (TPR). The AUC column chart (right) indicates the level of agreement between nucleotide reactivity and pairing status. A value of 0.5 indicates that nucleotide reactivity reports no base pairing information.

To further classify the PAN RNA secondary structure changes invoked by the METTL3 and FTO knockdowns, we performed classSNitch analysis (12, 56) (Fig. 5d). The classSNitch categorizes RNA structural changes based on the comparison of wild-type and experimental conditions by ranking eight features, e.g., Pearson correlation coefficient, contiguousness, change variance, etc., to determine local, global, or no changes to RNA structure (56). We compared PAN RNA reactivity profiles obtained in knockdown conditions to the wild-type reactivity profile, which served as a base input. The receiver operator curve (ROC, comparison between RNA classifier, wild-type, and knockdowns) generated from each profile was analyzed to estimate the accuracy of secondary structure prediction. The FTO knockdown was categorized as inducing local changes to the PAN RNA structure, with the ROC curve estimated area under the curve value of 0.86 (AUC, measurement of accuracy) (Fig. 5d). In contrast, the METTL3 knockdown was categorized as inducing the global changes to PAN RNA secondary structure, with the ROC curve estimated AUC value of 0.77 (Fig. 5d). AUC values ranging from 0.6-0.7 are regarded as having low accuracy. These results indicated that m^6^A can impact not only local PAN RNA structure but also its global folding, and as such, can hold substantial control over specific transcript’s biology. Also, the combination of genetic ablations with nucleotide-resolution RNA structure probing in vivo presents a potentially powerful method to dissect mechanisms governing a target RNA structure.

### PAN colocalizes with the m^6^A methylome components to varying degrees during the KSHV lytic replication

PAN RNA has been considered as a mainly nuclear lncRNA (87), with some reports suggesting that a fraction of the PAN transcript can leak to the cytoplasm to support specific intermolecular interactions (5, 12). Similar observations have been made for other predominantly nuclear lncRNAs, e.g., metastasis-associated lung adenocarcinoma transcript 1 (MALAT1) (88). We performed immunofluorescence analysis to assess the temporal and spatial subcellular distribution of our previously identified methylome enzymes, their potential for colocalization with PAN, and dependance on KSHV replication stage. We manually scored the immunofluorescence pattern of on average 10 cells from a number of microscopic fields and at each time point of infection, and found that PAN RNA was detectible as early as 8 hours post lytic induction at the immediate early stage of KSHV infection (T1). PAN RNA accumulated to the higest level at the early lytic stage (T2), and was located mainly within the nucleus (Fig. 6 and Supplementary Fig. 10).

**Fig. 6.**
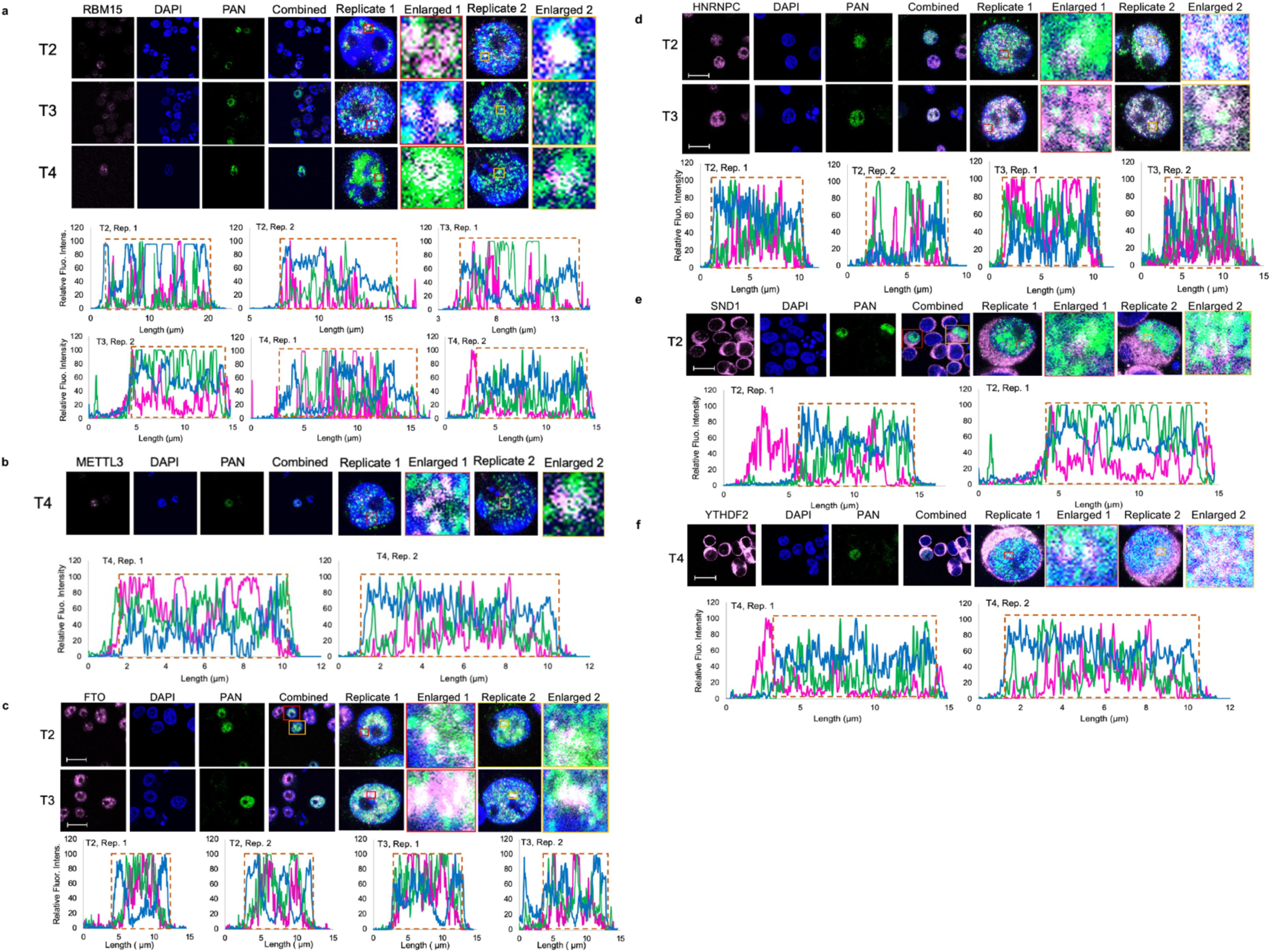
Confocal microscopy analysis of PAN and the indicated methylome components performed in BCBL-1 cells at latent (T0) and lytic (T1 - T4) stages of KSHV replication. The confocal microscopy micrographs are accompanied with related fluorescence intensity profiles for indicated magnified cells and identified interactions as follows: **a,** PAN-RBM15, **b**, PAN-METTL3, **c**, PAN-FTO, **d**, PAN-HNRNPC, **e**, PAN-SND1, and **f**. PAN-YTHDF2. DAPI staining of the nuclei is in blue, the methylome components are in pink and PAN RNA in green. (*n*=3). The fluorescence intensity profiles include lines color-coded according to staining; the red dashed line outlines the nucleus.

Recently, m^6^A-dependent colocalization of MALAT1 with RBM15 has been described, where RBM15 appeared to form small foci scattered over the nucleus and nuclear speckles (89). Our results are consistent with this observation, as we noted that RBM15 formed minute puncta that were contained within the nucleus. The PAN and RBM15 colocalization was most noticeable at the early (T2) and late lytic (T4) stages of viral lytic replication (Fig. 6a). Similarily, METTL3 formed a small foci within the subnuclear space (72, 90), and colocalization with PAN RNA was most evident during the late lytic stage of KSHV infection (T4) (Fig. 6b). We noted that the FTO localized abundantly to the nucleus (40, 91), and colocalized with PAN RNA to the highest degree during early (T2) and late lytic (T3) stages of viral replication (Fig. 6c). In the case of the m^6^A readers, HNRNPC resided abundantly within the nucleus, as previously reported (92), and colocalized with PAN to the highest degree at the early (T2) and late lytic (T3) stages of infection (Fig. 6d). However, both SND1 and YTHDF2 were found to have profuse cytoplasmic localization, with small fractions of both enzymes being subnuclear. In particular, PAN and SND1 colocalization was detected during the early lytic stage (T2) in the nucleus (Fig. 6e), while PAN and YTHDF2 colocalization was noted at the late lytic stage of KSHV infection (T4), and also occurred within the nucleus (Fig. 6f). These data highlight that the progression of KSHV replication does not influence the localization of the m^6^A methylome components identified to influence PAN RNA modification, and that all of them are temporarily and spatially available to regulate or mediate the phenotypic effect of PAN m^6^A.

## DISCUSSION

The intricate and interconnected nature of the KSHV transcriptome bestows the expression of numerous long non-coding RNAs, which regulate viral and cellular gene expression by means that are not fully understood (11, 43, 44, 90). These non-coding transcripts themselves endure regulation by various mechanisms, including epitranscriptomic modifications. Recent studies have revealed the critical impact of m^6^A residues on the progression of KSHV latent and lytic stages of replication (88, 91). No studies, however, addressed the dynamics of the m^6^A landscape and its abundance in the context of specific non-coding viral transcript, and during the progression of viral replication.

In our study, we provide the first comprehensive insight into the m^6^A status of KSHV-encoded PAN RNA, the key regulator of viral lytic reactivation (7, 92, 93). Using two independent mapping approaches, i.e., the 4SedTTP RT with next-generation sequencing (Fig. 1, Supplementary Fig. 1b and 2) and direct RNA sequencing (Fig. 2), we revealed that PAN RNA can carry up to 5 m^6^A sites. These modifications localize within PAN RNA secondary structure motifs that we have previously shown to display low SHAPE and low Shannon entropy (12), which are characteristic of biologically significant RNA motifs (94–97). We noted that PAN m^6^A status is dynamic and changes during the progression of KSHV replication, achieving the highest level during the late lytic stages. This process might operate as a fine-tuning mechanism, coordinating the function and biology of PAN RNA, as it has been demonstrated for other transcripts (30, 31, 33, 98). We established the PAN m^6^A consensus sequence at the A/G/Cm^6^ACA/U/C, and noted that the motif shows higher sequence variability than the motif identified for cellular lncRNAs (79, 99, 100) (Supplementary Fig. 4). Using a newly developed method that we termed <u>S</u>elenium-modified deoxythymidine triphosphates (SedTTP) RT and <u>L</u>igation <u>A</u>ssisted <u>P</u>CR analysis (SLAP), we gained insight into the modification frequency at identified m^6^A sites. We reasoned that the quantification of the extent of modification could advance the understanding of how changes in PAN modification pattern, the site and fraction, are modulated during the KSHV infection. This is also a crucial parameter for investigating the biological significance of PAN m^6^A. For example, it has been shown that only one out of fourteen m^6^A detected on HOX antisense intergenic lncRNA (HOTAIR), which displayed the most consistent modification, plays a critical role in HOTAIR-mediated cellular proliferation, colony formation, and HOTAIR nuclear retention (22). Also, one out of four m^6^A detected in MALAT1, which shows over 80% modification frequency in two cell lines (101), is attributed to the m^6^A-induced structural change associated with m^6^A reader binding (100). In our studies, the measurement of the m^6^A frequency at each site identified was the highest during the late lytic stages of KSHV infection (Fig. 3).

We also tackled the identification of methylome components that regulate PAN m^6^A status. Using a combination of proteomic approaches (Fig. 4) and confocal microscopy analysis (Fig. 6), we showed that the writer METTL3, writer recruiter RBM15, and eraser FTO, are not only temporarily and spatially available for PAN modification but also directly interact with the target RNA. Knockdown of any of these enzymes diminished PAN RNA expression at various stages of KSHV lytic replication (Fig. 7e) and abolished the m^6^A modification frequency (Fig. 4 and Supplementary Fig. 8). We also identified m^6^A readers that directly associate with PAN RNA, i.e., SND1, HNRNPC, and YTHDF2, (Fig. 4). SND1 has been reported to bind and stabilize m^6^A modified viral ORF50 mRNA, which is essential for the KSHV replication (37). That study also identified the SND1-binding motif as a U-tract immediately followed by the Gm^6^AC motif. We found similar motif near m^6^A at position 672 on PAN RNA. Notably, U-rich motifs found adjacent to m^6^A are also bound by RBM15 and HNRNPC (79, 102), both of which, we found to associate with PAN RNA (Fig. 4). Another PAN m^6^A reader, the YTHDF2, has been shown to recruit the CCR4-NOT deadenylase complex and promote the degradation of modified transcripts (103). The knockdown of YTHDF2 has been shown to trigger elevated expression of KSHV transcripts and release of virions, which is suggestive of YTHDF2 functioning as a restriction factor for KSHV lytic replication (104). We showed that YTHDF2 directly associates with PAN RNA at the late lytic stage of KSHV infection (Fig. 4 and Fig. 6), during which our target transcript is the most extensively modified (Fig. 1 and 3). Thus, it is feasible that the YTHDF2 association with PAN might be involved in the regulation of PAN RNA turnover (98, 105, 106).

We also delineated the structural effect of m^6^A in the context of full-length PAN RNA. The m^6^A modification is known to diminish the ability of RNA to form duplexes, favoring the linear, un-paired form of RNA. This reduction in structure provides greater access to RNA by RNA-binding proteins that associate with single-stranded motifs (15, 102, 107). Another study has shown that the m^6^A-directed structural effect strongly depends on the modified RNA’s structural context, and when neighboring by a 5ʹ bulge, m^6^A stabilizes rather than disrupts base pairing (108). Our study showed that the knockdown of METTL3 or FTO results in the alternation of the local and global secondary structure of PAN RNA (Fig. 5). The most striking structural change was invoked by METTL3 ablation, as the stabilizing ENE motif, known to be involved in the triple helix formation, was found to be disrupted (83, 84). We noted that the ENE U-rich loop residues (nts 901 - 905 and 947 - 953) were separated, and engaged in the formation of two distinct hairpins, making them unlikely to sequester PAN poly(A) tail during the triple helix formation. These results suggests that the m^6^A modification holds powerful grip over RNA structure, that can ether result in the disruption of functionally critical RNA motifs or unmasking new ones, which in turn, can influence overall RNA biology, e.g., cellular fate, stability, interactions with binding partners. Proteins binding also induces changes to RNA structure, and since m^6^A can regulate RNA structural access to their respective protein partners, it remains to be determined whether the observed structural changes are due to the altered m^6^A modification pattern or RNA-protein interactions.

This work represents the most comprehensive overview of the dynamic interplay that takes place among the cellular epitranscriptomic machinery and a specific viral long non-coding RNA. Our findings provide thought-provoking clues regarding the influence of m^6^A modification over PAN RNA secondary structure and biology, e.g., RNA stability and turnover. Only recently has it been shown that the PAN expression is upregulated in cells with the knockdown of nonsense-mediated decay (NMD) factors (109). Considering that our data show PAN association with YTHDF2, which is known to interact with the NMD complex and trigger m^6^A-mediated RNA decay (98), and that METTL3 and FTO knockdowns affect PAN expression levels and the secondary structure of its critical stabilizing motif, ENE, the m^6^A modification can hold clues to the conundrum of PAN RNA turnover.

## AVAILABILITY

Data supporting reported results can be found at NCBI Sequence Read Archive (SRA) submission numbers SUB9396779, SUB9396663, and SUB9421156. All data are reported in the text and supplementary materials. At home-developed R scripts are available at https://github.com/jzs0165.

## ACCESSION NUMBERS

The PAN RNA sequence can be found at the accession number: U50139.1.

## SUPPLEMENTARY DATA

Supplementary Data are available at NAR online.

## ACKNOWLEDGEMENT

The authors would like to thank Dr. Denise Whitby (NIH/NCI-Frederick) for BCBL-1 cell line, Dr. Jennifer Miller (NIH/NCI-Frederick) for constructive criticism of the manuscript, Damian Waits (Auburn University), Philip Forrest, and Chris Solinski (COSAM IT, Auburn University) for their bioinformatic and technical support.

## SUPPLEMENTARY DATA TABLES AND FIGURES

**Supplementary Fig. 1 a**, The KSHV infectivity cycle. Latent (T0) and lytic stages of KSHV infectivity cycle are indicated. Lytic stage is further divided into the following time points: T1 – immediate early (8h post induction, h pi), T2 – early (24 h pi), T3 (48 h pi) and T4 late lytic (72 h pi) based on distinct gene expression panel. PAN RNA expression is the most abundant at early lytic stage (T2). **b**, The schematic overview of 4SedTTP RT and next-generation sequencing methodology. Total RNA was subjected to the K7 mRNA depletion with a K7-specific biotinylated antisense probe (green). Fractions depleted of K7 mRNA were then subjected to the affinity capture of PAN with specific biotinylated antisense probes (yellow). Captured PAN RNA (red) was reverse transcribed in the presence of 4SedTTP to induce RT stops at m^6^A (bright green). The 4SedTTP RT experimental and control reactions were subjected to library preparation and next-generation sequencing.

**Supplementary Fig. 2** 4SedTTP RT strategy for detection of m^6^A. **a**, The electropherogram representing the results of initial assessment of 4SedTTP RT strategy. Fluorescent primer (17 nt) and in vitro transcript (40 nt) carrying one m^6^A residue were used in the RT reaction in the presence of either 4SedTTP (+) or dTTP (-). The presence of 4SedTTP in RT reactions induced the truncation of cDNA products (27 nt) as compared to the full-length cDNA product (40 nt). **b**, The plot represents next-generation sequencing analysis performed on in vitro synthesized transcripts: one carrying m^6^A at positions 13 and 82 (blue), and another unmodified transcript. The X-axis represents nucleotide sequence, the Y-axis indicates m^6^A_FC_ values.

**Supplementary Fig. 3** Direct PAN RNA sequencing analysis. **a**, Ternary diagrams depicting the mismatch directionality for m^6^A residues in (from left to right) 0% modified, 100% modified control transcripts and PAN RNA expressed at latent (T0) and lytic (T3) stages of KSHV infection. The scale on triangle sides reflects base proportions. **b**, Total insertions and deletions (INDEL), **c**, Base-quality (BQ), **d**, Normalized current intensity values assessed for PAN RNA expressed during the latent (blue) and lytic (orange) stages of KSHV infection and contrasted with values obtained for 100% modified (yellow) and unmodified transcripts (gray). **e**, Predicted stoichiometry of m^6^A modifications using k-nearest neighbors (KNN) algorithm that classifies reads as modified or non-modified. X-axis represents all adenosines in 100% and 0% modified control PAN transcripts, and specific modified adenosines on PAN RNA expressed during latency (T0, blue), and lytic KSHV replication (T3, orange). Y-axis represents m^6^A recovery percentage.

**Supplementary Fig. 4** The m^6^A consensus on PAN RNA. **a**, The list of sequences carrying m6A within PAN and selected cellular lncRNAs, i.e., Growth Arrest Specific 5 (GAS5, Ni et al., 2019), HOX antisense intergenic RNA (HOTAIR, Porman et al., 2020), Metastasis Associated Lung Adenocarcinoma Transcript 1(MALAT1, Zhou et al., 2016; Wang et al., 2021), and X-inactive specific transcript (XIST, Patil et al., 2016). **b**, MEME analysis of PAN RNA m^6^A signatures. c, MEME analysis of the m^6^A sequence environment in cellular lncRNAs.

**Supplemental Fig. 5** The RAP MS analysis for PAN RNA m^6^A methylome. **a**, Normalized expression of PAN RNA assessed by RT qPCR of total RNA samples extracted during KSHV latent (T0) and lytic (T3) stages of replication, both with and without PAN RNA-protein crosslinking and affinity capture. **b**, The lncRNA-protein interactions network (LPI) for the PAN RNA m^6^A methylome visualized by Cytoscape. Proteins that were found to associate with PAN RNA are shown in purple (writers), blue (readers), and green (erasers).

**Supplemental Fig. 6** The normalized expression of methylome-associated mRNAs encoding RBM15, METTL3, FTO, HNRNPC, YTHDF2, SND1 during the latent and lytic stages of KSHV replication. MALAT1 RNA was used as endogenous control RNA (n=3).

**Supplementary Fig. 7** The knockdowns of m^6^A methylome components affect PAN RNA expression levels. The normalized mRNAs expression (Y-axis) of **a**, RBM15, **b**, METTL3, **c**, FTO in BCBL-1 knockdowns (KD) cell lines. The X-axis corresponds to the latent (T0) and lytic (T1-T4) stages of KSHV infection. Control samples represent latent BCBL-1 cells treated with scrambled siRNAs. **d**, Western blots representing the efficiency of knockdown on the level of proteins expression. Tubulin was used as a control. e, PAN RNA expression in knockdown cell lines versus control at specified time points of KSHV replication. MALAT1 was used for normalization (n = 4).

**Supplementary Fig. 8 a**, The graphs represent m^6^A_RATIO_ (Y-axis) against PAN RNA sequence (X-axis). The peaks correspond to the expected RT stop 1 nt upstream of m^6^A. The threshold line (red) for calling m^6^A peaks is set at 0.2 and it was calculated by dividing the average m^6^A_RATIO_ values calculated for all PAN nucleotides divided by the total number of nucleotides. **b**, The charts representing the signal-to-noise ratio for PAN sequence overlapping m^6^A within specified nucleotide window (X-axis) in BCBL-1 for RBM15, METTL3, and FTO knockdown (KD) cell lines. Time points of KSHV infection are color-coded as follows: latency - T0 in blue; late lytic stage - T3 in yellow.

**Supplementary Fig. 9** Step-plots representing reactivity profiles for PAN RNA probed in **a**, wild-type, **b**, METTL3. **c**, FTO knockdown BCBL-1 cells. X-axis corresponds to PAN RNA nucleotide sequence, Y-axis represents SHAPE-MaP reactivity values. Nucleotides 1-14 and 1056-1077 were not covered in the analysis due to primer annealing. The position of the expression and nuclear retention (ENE) motif is indicated with green square.

**Supplementary Fig. 10** Confocal fluorescent microscopy analysis of PAN and related m^6^A methylome components performed in BCBL-1 cells during the latent (T0) and lytic (T1 – T4) stages of KSHV replication. The confocal microscopy micrographs representing the individual methylome enzymes (in pink) are ordered as follows: **a**, RBM15, **b**, METTL3, **c**, FTO, **d**, HNRNPC, **e**, SND1, and **f**, YTHDF2. DAPI staining of the nuclei is in blue, and PAN RNA is green.

