## Supplemental Files for "The m^6^A landscape of polyadenylated nuclear (PAN) RNA and its related methylome in the context of KSHV replication"

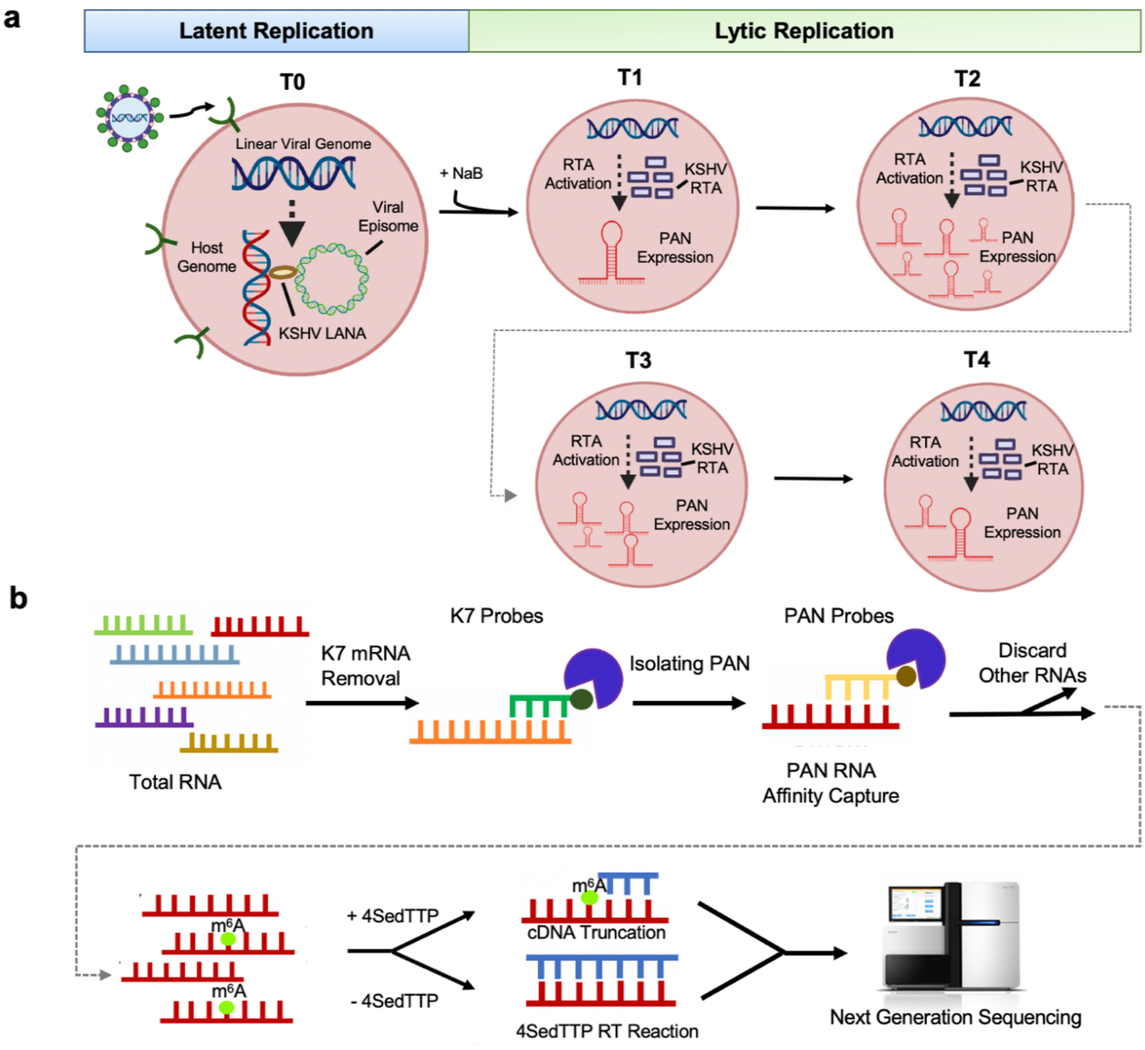


**Supplementary Fig. 1 a, The KSHV infectivity cycle**. Latent (T0) and lytic stages of KSHV infectivity cycle are indicated. Lytic stage is further divided into the following time points: T1 – immediate early (8h post induction, h pi), T2 – early (24 h pi), T3 (48 h pi) and T4 late lytic (72 h pi) based on distinct gene expression panel. PAN RNA expression is the most abundant at early lytic stage (T2). **b, The schematic overview of 4SedTTP RT and next-generation sequencing methodology.** Total RNA was subjected to the K7 mRNA depletion with a K7-specific biotinylated antisense probe (green). Fractions depleted of K7 mRNA were then subjected to the affinity capture of PAN with specific biotinylated antisense probes (yellow). Captured PAN RNA (red) was reverse transcribed in the presence of 4SedTTP to induce RT stops at $m^6A$ (bright green). The 4SedTTP RT experimental and control reactions were subjected to library preparation and next-generation sequencing.

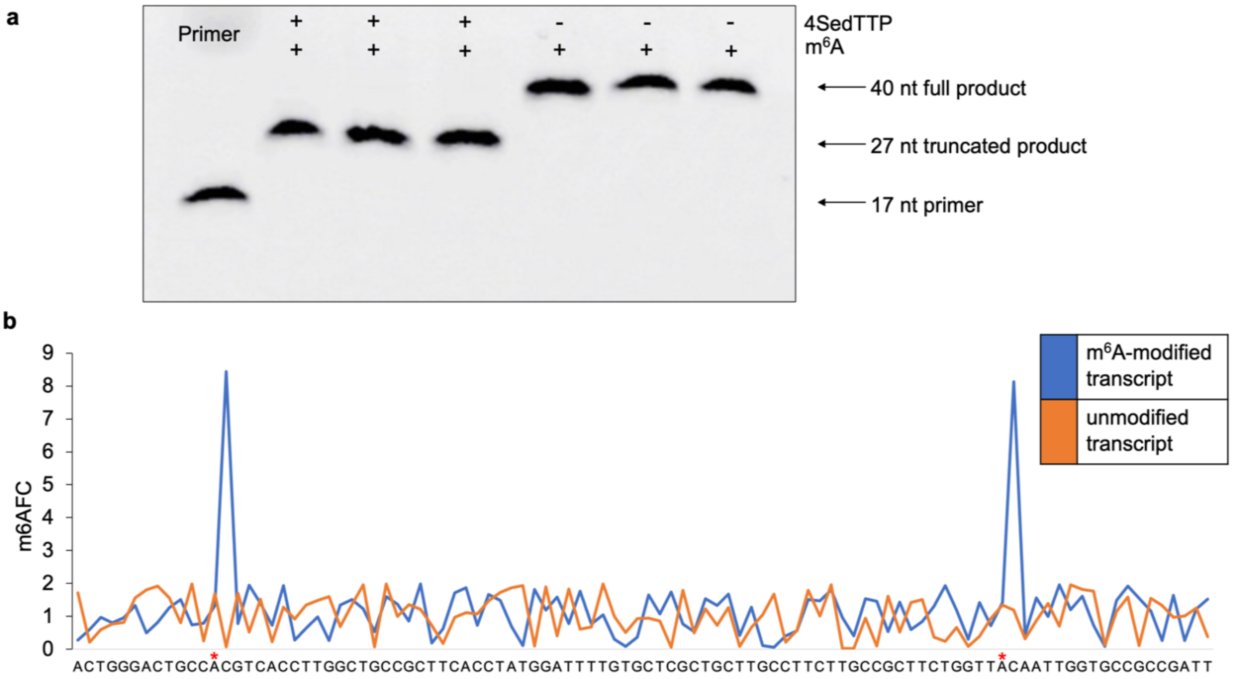


**Supplementary Fig. 2 4SedTTP RT strategy for detection of m$^6$A. a**, The electropherogram representing the results of initial assessment of 4SedTTP RT strategy. Fluorescent primer (17 nt) and in vitro transcript (40 nt) carrying one m$^6$A residue were used in the RT reaction in the presence of either 4SedTTP (+) or dTTP (-). The presence of 4SedTTP in RT reactions induced the truncation of cDNA products (27 nt) as compared to the full-length cDNA product (40 nt). **b**, The plot represents next-generation sequencing analysis performed on in vitro synthesized transcripts: one carrying m$^6$A at positions 13 and 82 (blue), and another unmodified transcript. The X-axis represents nucleotide sequence, the Y-axis indicates m$^6$A$_{FC}$ values.

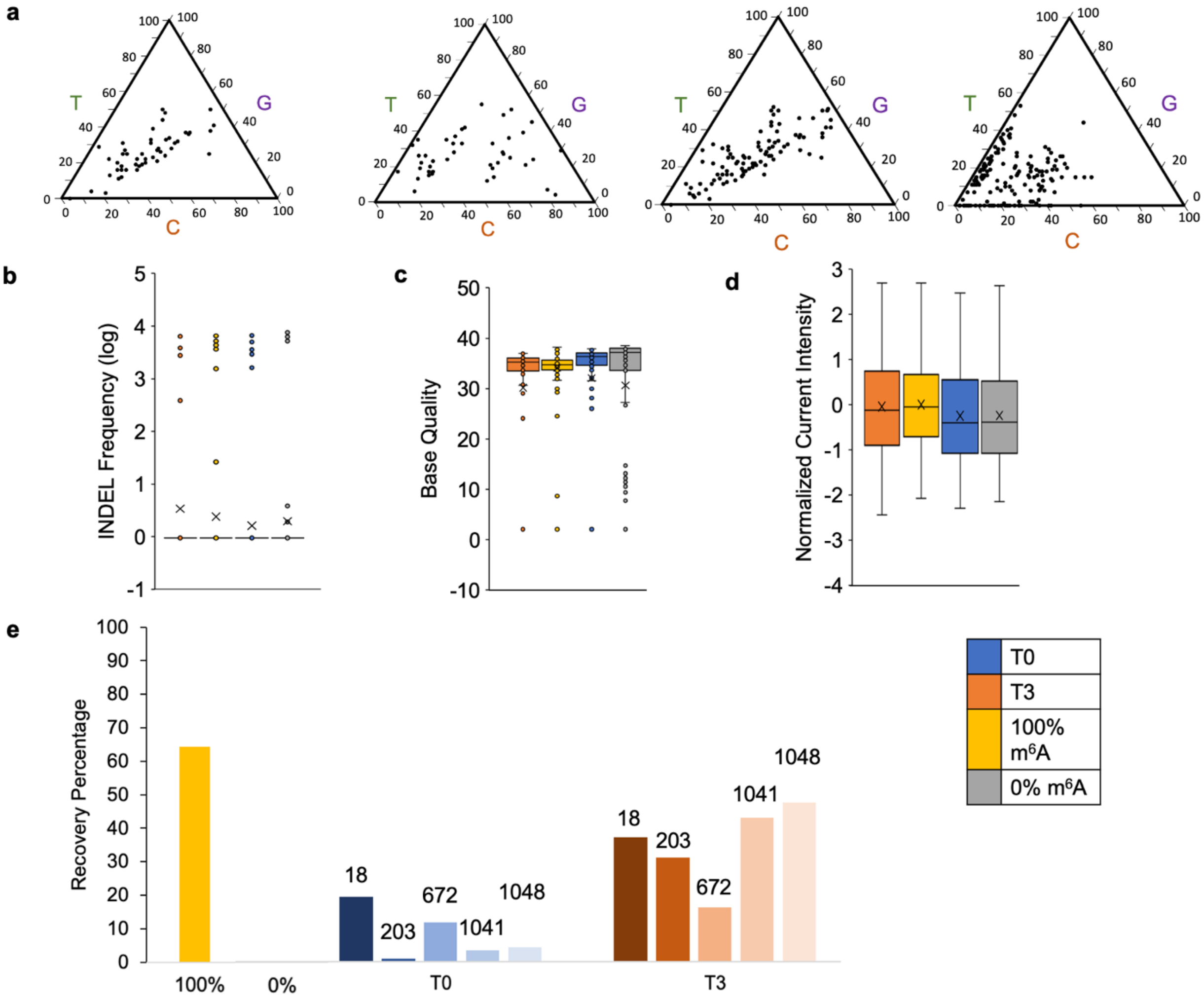


**Supplementary Fig. 3 Direct PAN RNA sequencing analysis. a**, Ternary diagrams depicting the mismatch directionality for $m^6A$ residues in (from left to right) 0% modified, 100% modified control transcripts and PAN RNA expressed at latent (T0) and lytic (T3) stages of KSHV infection. The scale on triangle sides reflects base proportions. **b**, Total insertions and deletions (INDEL), **c**, base-quality (BQ), **d**, normalized current intensity values assessed for PAN RNA expressed during the latent (blue) and lytic (orange) stages of KSHV infection and contrasted with values obtained for 100% modified (yellow) and unmodified transcripts (gray). **e**, Predicted stoichiometry of $m^6A$ modifications using k-nearest neighbors (KNN) algorithm that classifies reads as modified or non-modified. X-axis represents all adenosines in 100% and 0% modified control PAN transcripts, and specific modified adenosines on PAN RNA expressed during latency (T0, blue), and lytic KSHV replication (T3, orange). Y-axis represents $m^6A$ recovery percentage.

a

| lncRNA | $m^6A$ site | Motif |
|---|---|---|
| PAN RNA | 18 | UC$m^6$ACC |
| | 203 | AA$m^6$ACA |
| | 672 | UG$m^6$ACA |
| | 1041 | CA$m^6$ACU |
| | 1048 | CA$m^6$ACA |
| GAS5 | 362 | GG$m^6$ACU |
| HOTAIR | 783 | GA$m^6$ACG |
| MALAT1 | 2515 | GG$m^6$ACU |
| | 2577 | GG$m^6$ACU |
| | 2611 | GG$m^6$ACU |
| | 2674 | AG$m^6$ACA |
| | 2684 | GA$m^6$ACA |
| | 2698 | GA$m^6$ACC |
| | 2720 | GG$m^6$ACU |
| XIST* | | AA$m^6$ACA |
| | | UG$m^6$ACU |
| | | AC$m^6$ACA |

b

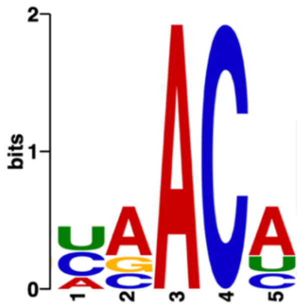


c

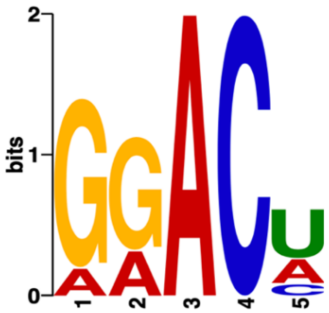


**Supplementary Fig. 4 The $m^6$A consensus on PAN RNA. a**, The list of sequences carrying $m^6$A within PAN and selected cellular lncRNAs, i.e., Growth Arrest Specific 5 (GAS5, *Ni et al., 2019*), HOX antisense intergenic RNA (HOTAIR, *Porman et al., 2020*), Metastasis Associated Lung Adenocarcinoma Transcript 1(MALAT1, *Zhou et al., 2016; Wang et al., 2021*), and X-inactive specific transcript (XIST, *Patil et al., 2016*). **b**, MEME analysis of PAN RNA $m^6$A signatures. **c**, MEME analysis of the $m^6$A sequence environment in cellular lncRNAs.

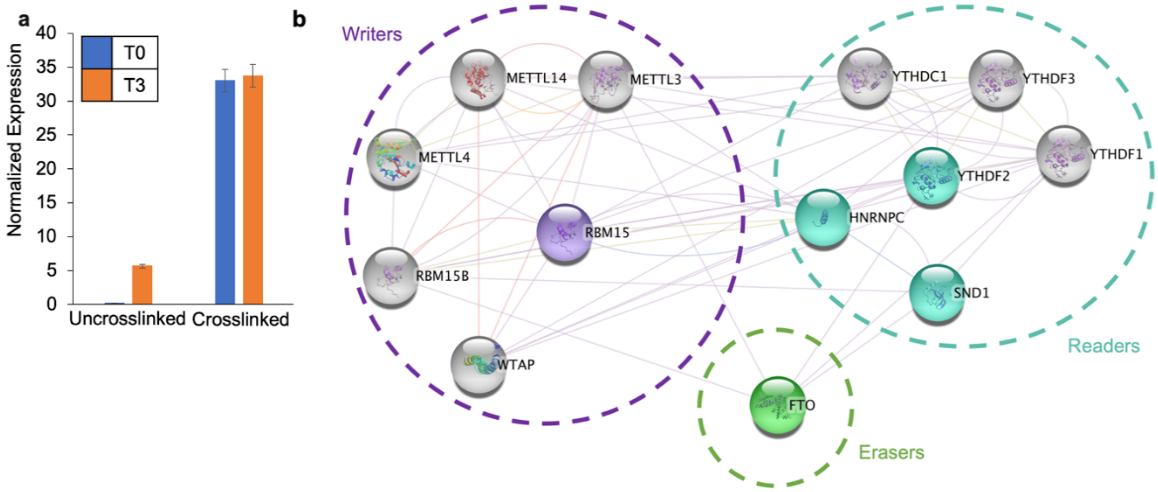


**Supplemental Fig. 5 The RAP MS analysis for PAN RNA m$^6$A methylome. a**, Normalized expression of PAN RNA assessed by RT qPCR of total RNA samples extracted during KSHV latent (T0) and lytic (T3) stages of replication, both with and without PAN RNA-protein crosslinking and affinity capture. **b**, The lncRNA-protein interactions network (LPI) for the PAN RNA m$^6$A methylome visualized by Cytoscape. Proteins that were found to associate with PAN RNA are shown in purple (writers), blue (readers), and green (erasers).

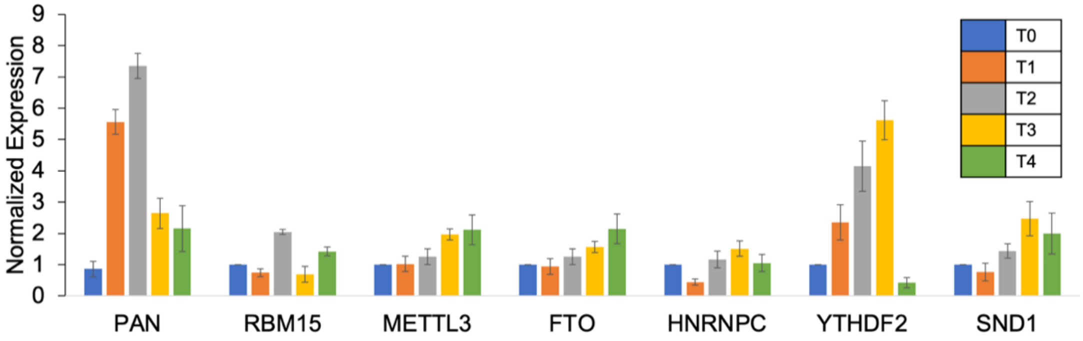


**Supplemental Fig. 6** The normalized expression of methylome-associated mRNAs encoding RBM15, METTL3, FTO, HNRNPC, YTHDF2, SND1 during the latent and lytic stages of KSHV replication. MALAT1 RNA was used as endogenous control RNA (n=3).

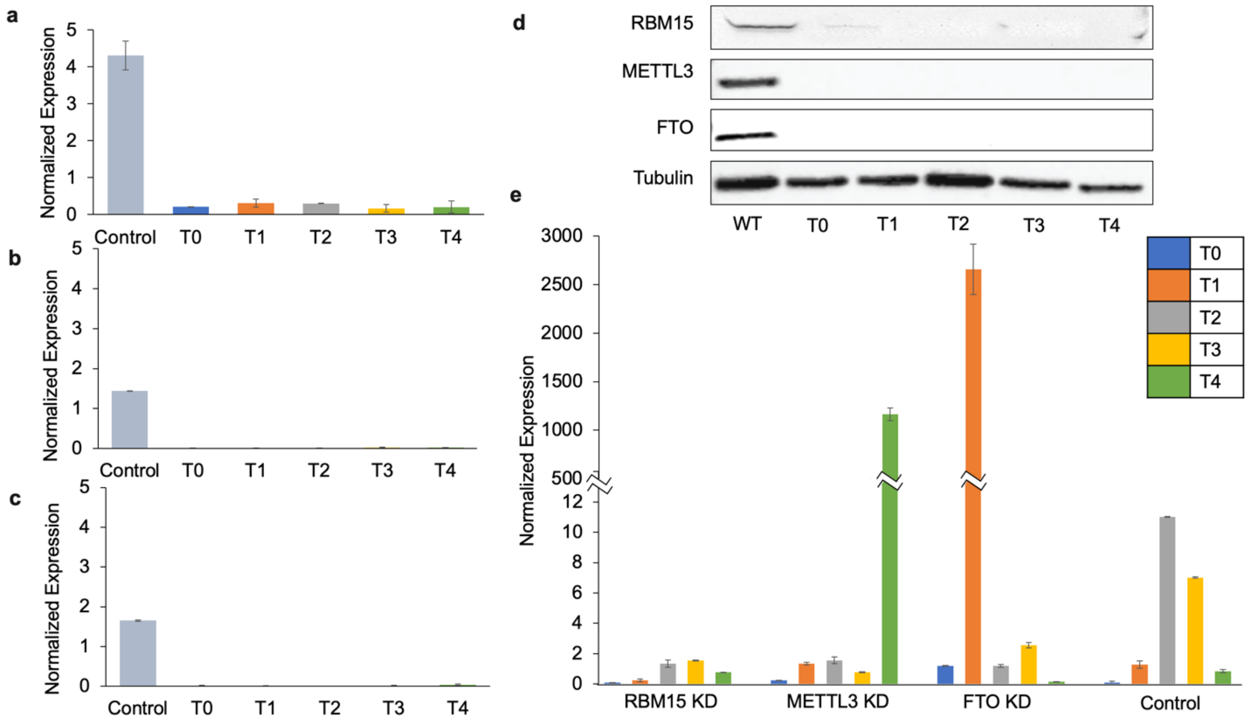


**Supplementary Fig. 7 The knockdowns of m$^6$A methylome components affect PAN RNA expression levels.** The normalized mRNAs expression (Y-axis) of **a**, RBM15, **b**, METTL3, **c**, FTO in BCBL-1 knockdowns (KD) cell lines. The X- axis corresponds to the latent (T0) and lytic (T1-T4) stages of KSHV infection. Control samples represent latent BCBL-1 cells treated with scrambled siRNAs. **d**, Western blots representing the efficiency of knockdown on the level of proteins expression. Tubulin was used as a control. **e**, PAN RNA expression in knockdown cell lines versus control at specified time points of KSHV replication. MALAT1 was used for normalization (n = 4).

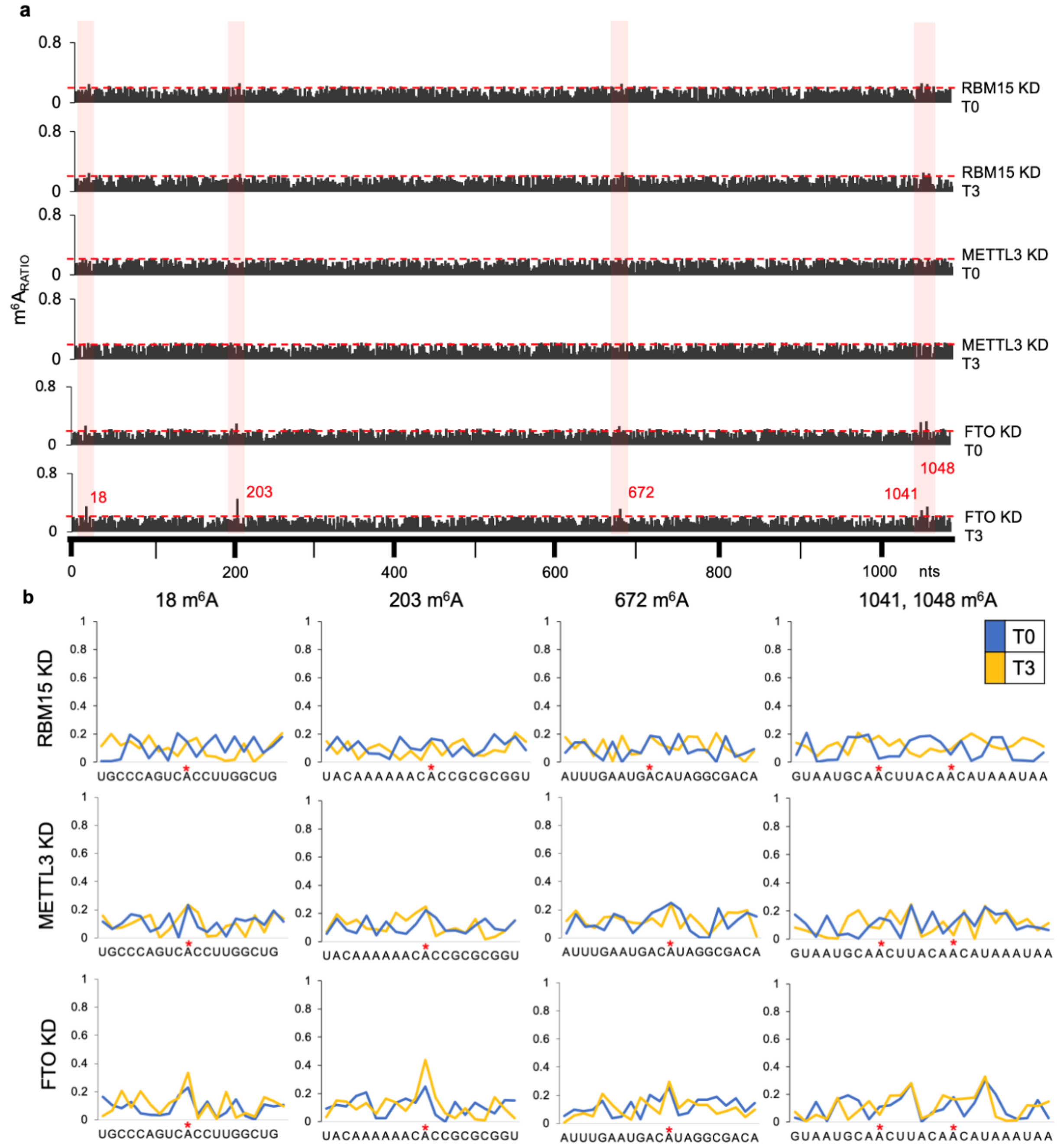


**Supplementary Fig. 8 a,** The graphs represent m$^6$A$_{RATIO}$ (Y-axis) against PAN RNA sequence (X-axis). The peaks correspond to the expected RT stop 1 nt upstream of m$^6$A. The threshold line (red) for calling m$^6$A peaks is set at 0.2 and it was calculated by dividing the average m$^6$A$_{RATIO}$ values calculated for all PAN nucleotides divided by the total number of nucleotides. **b,** The charts representing the signal-to-noise ratio for PAN sequence overlapping m$^6$A within specified nucleotide window (X-axis) in BCBL-1 for RBM15, METTL3, and FTO knockdown (KD) cell lines. Time points of KSHV infection are color-coded as follows: latency - T0 in blue; late lytic stage - T3 in yellow.

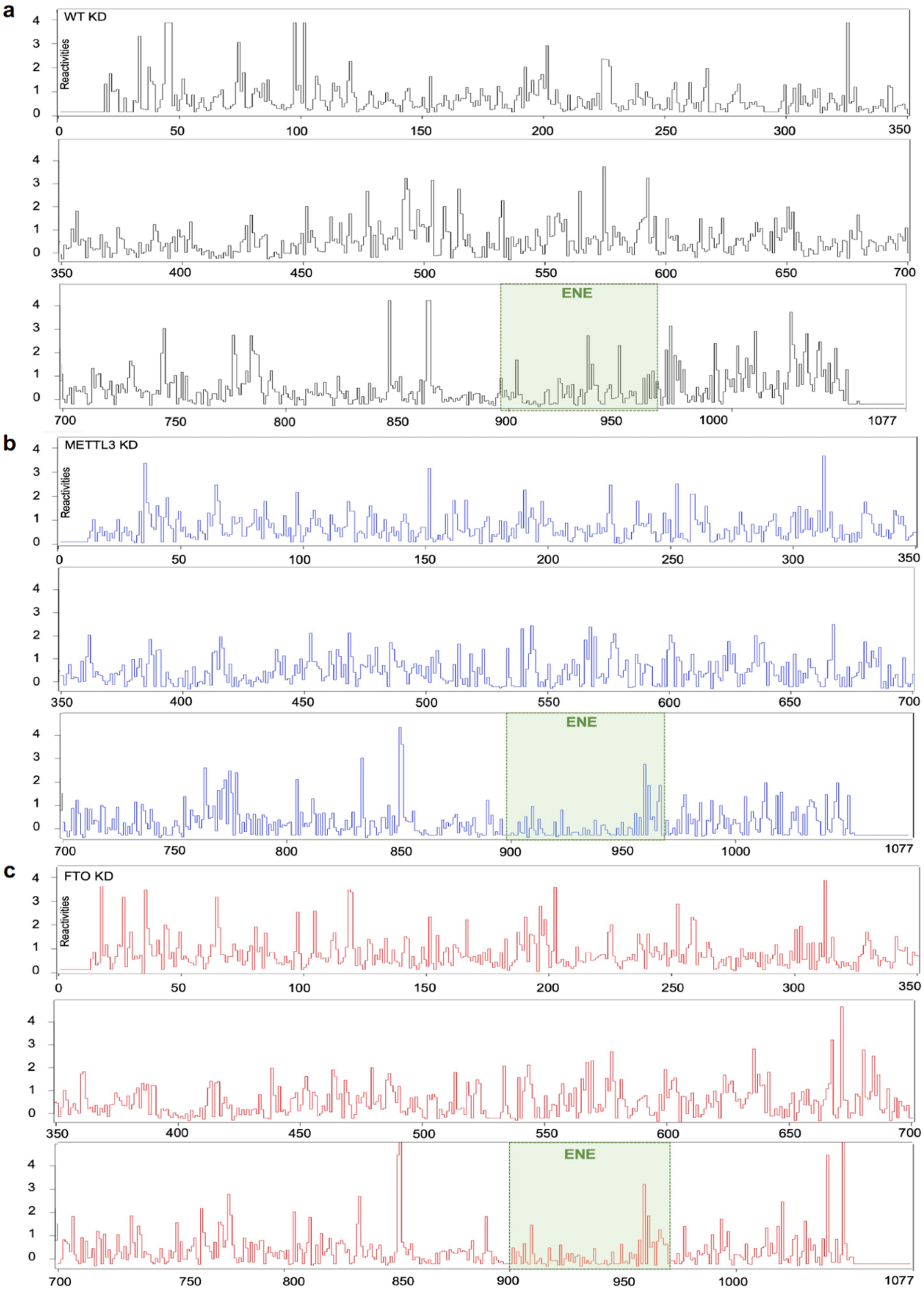


**Supplementary Fig. 9** Step-plots representing reactivity profiles for PAN RNA probed in **a,** wild-type, **b,** METTL3. **c,** FTO knockdown BCBL-1 cells. X- axis corresponds to PAN RNA nucleotide sequence, Y-axis represents SHAPE-MaP reactivity values. Nucleotides 1-14 and 1056-1077 were not covered in the analysis due to primer annealing. The position of the expression and nuclear retention (ENE) motif is indicated with green square.

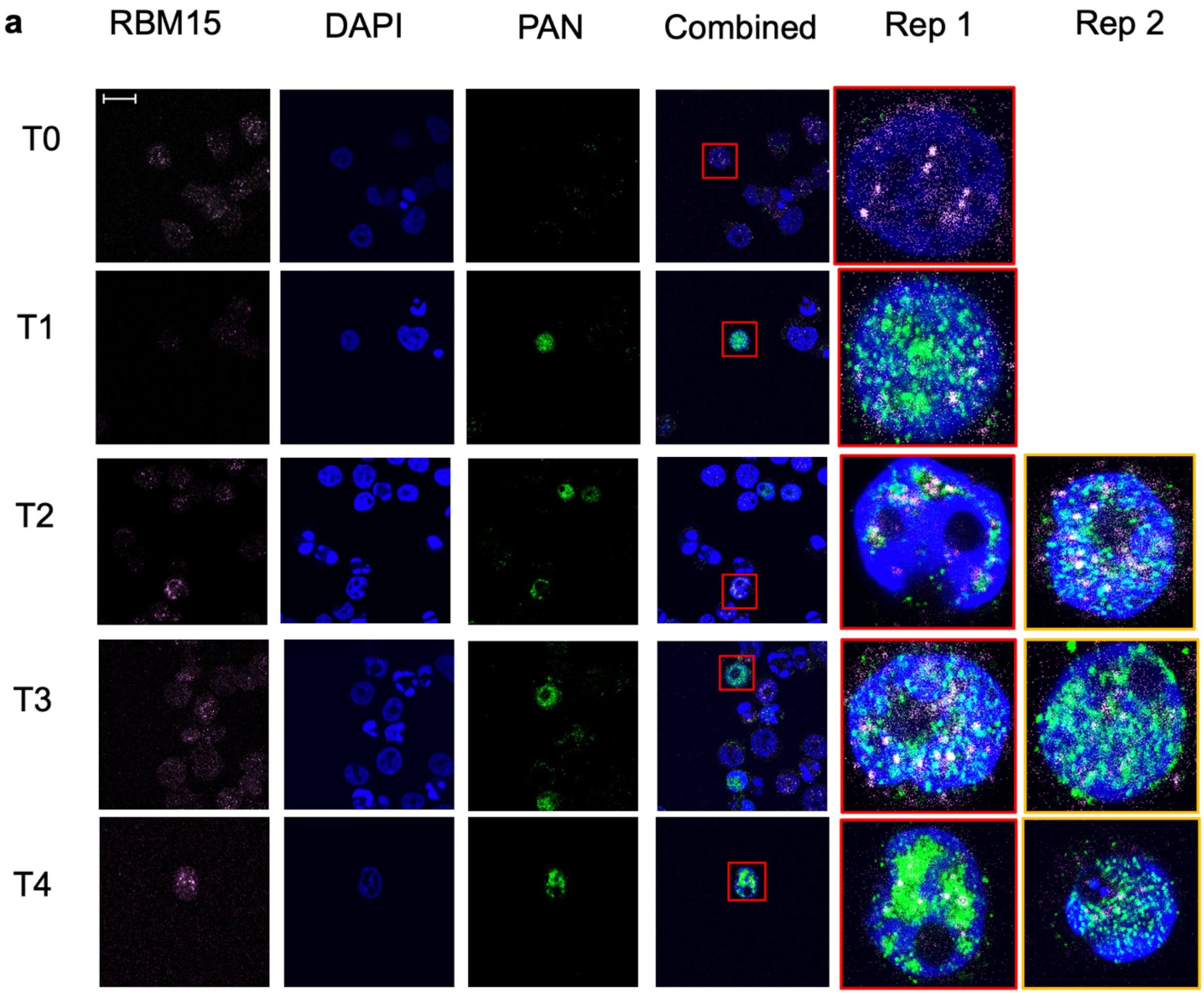
a
RBM15
DAPI
PAN
Combined
Rep 1
Rep 2
T0
T1
T2
T3
T4

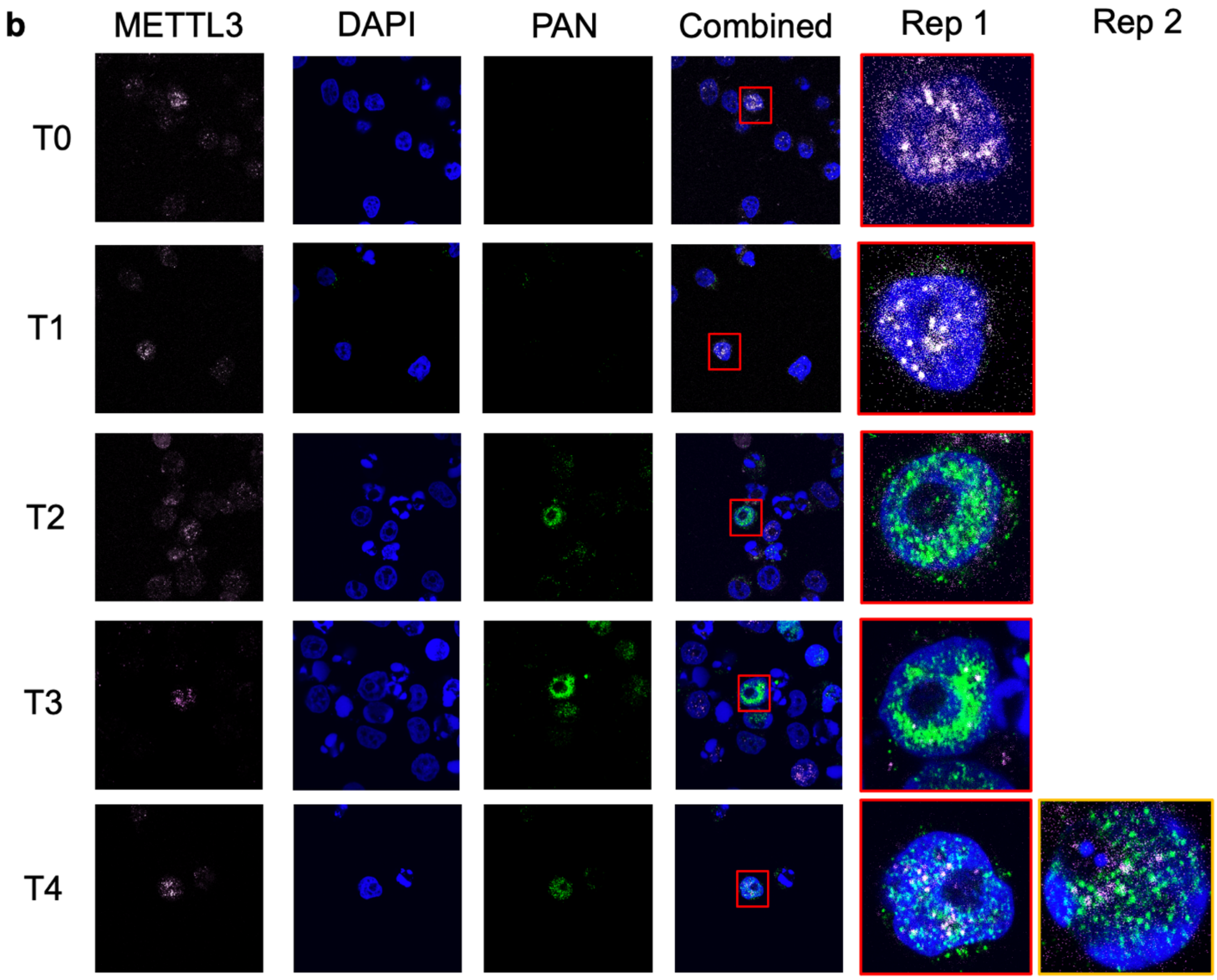

b
METTL3
DAPI
PAN
Combined
Rep 1
Rep 2
T0
T1
T2
T3
T4

c

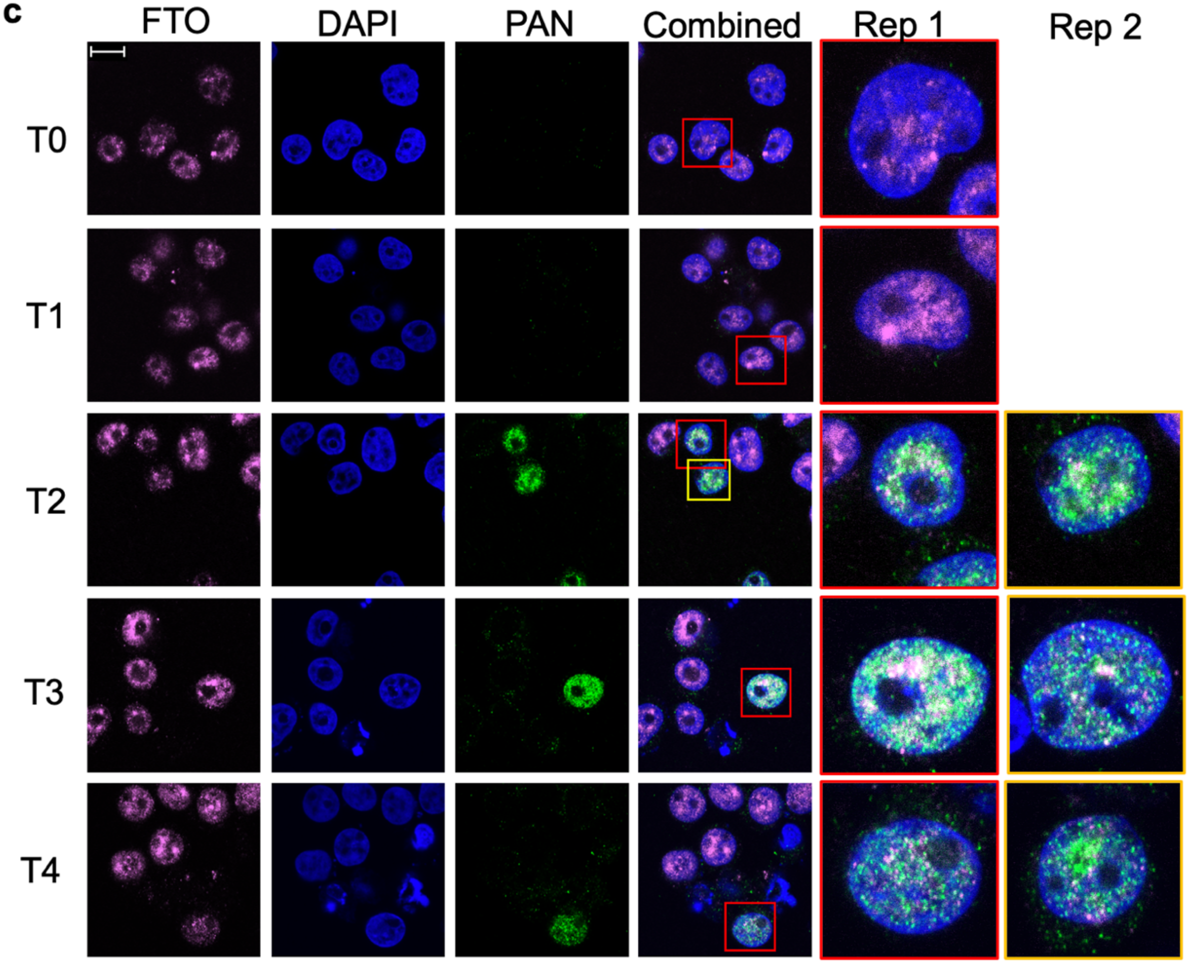

**d**

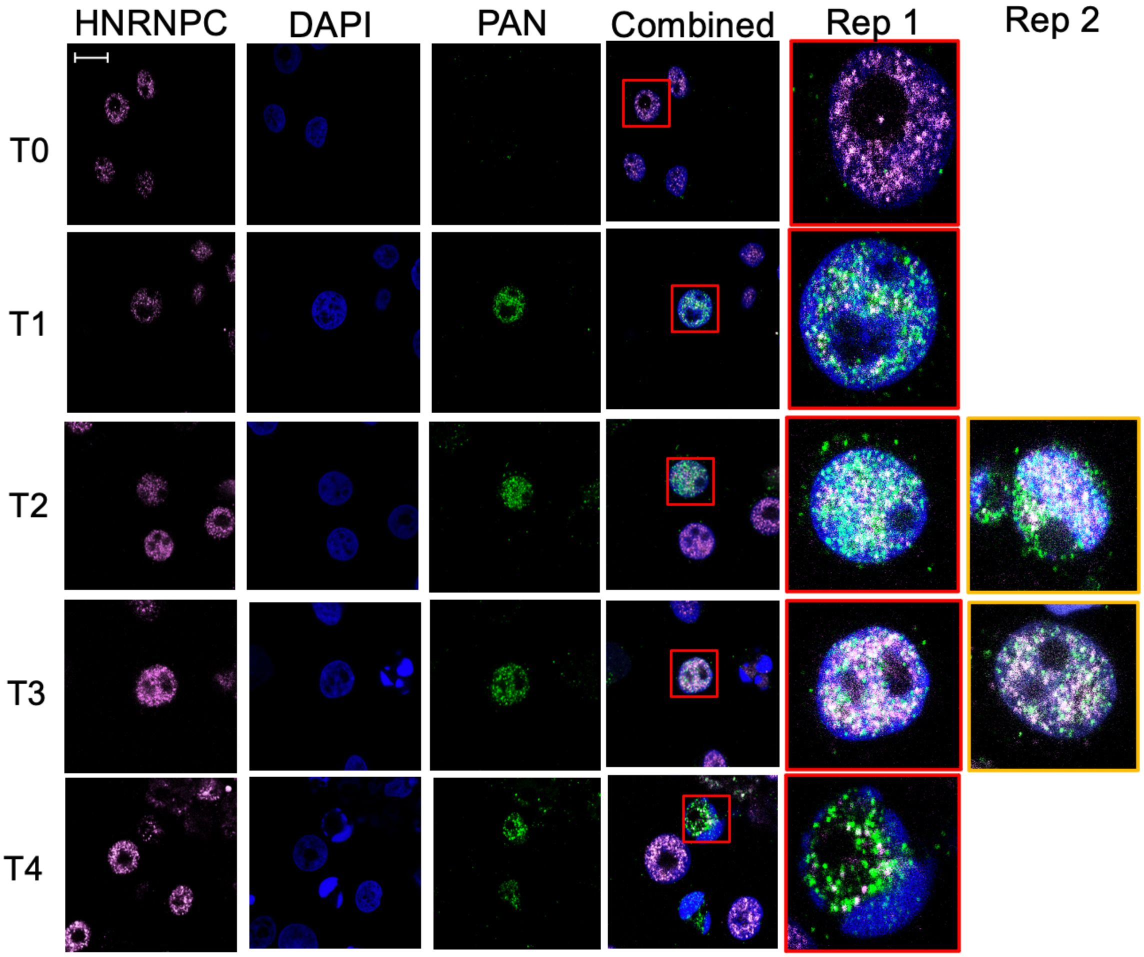

**e**

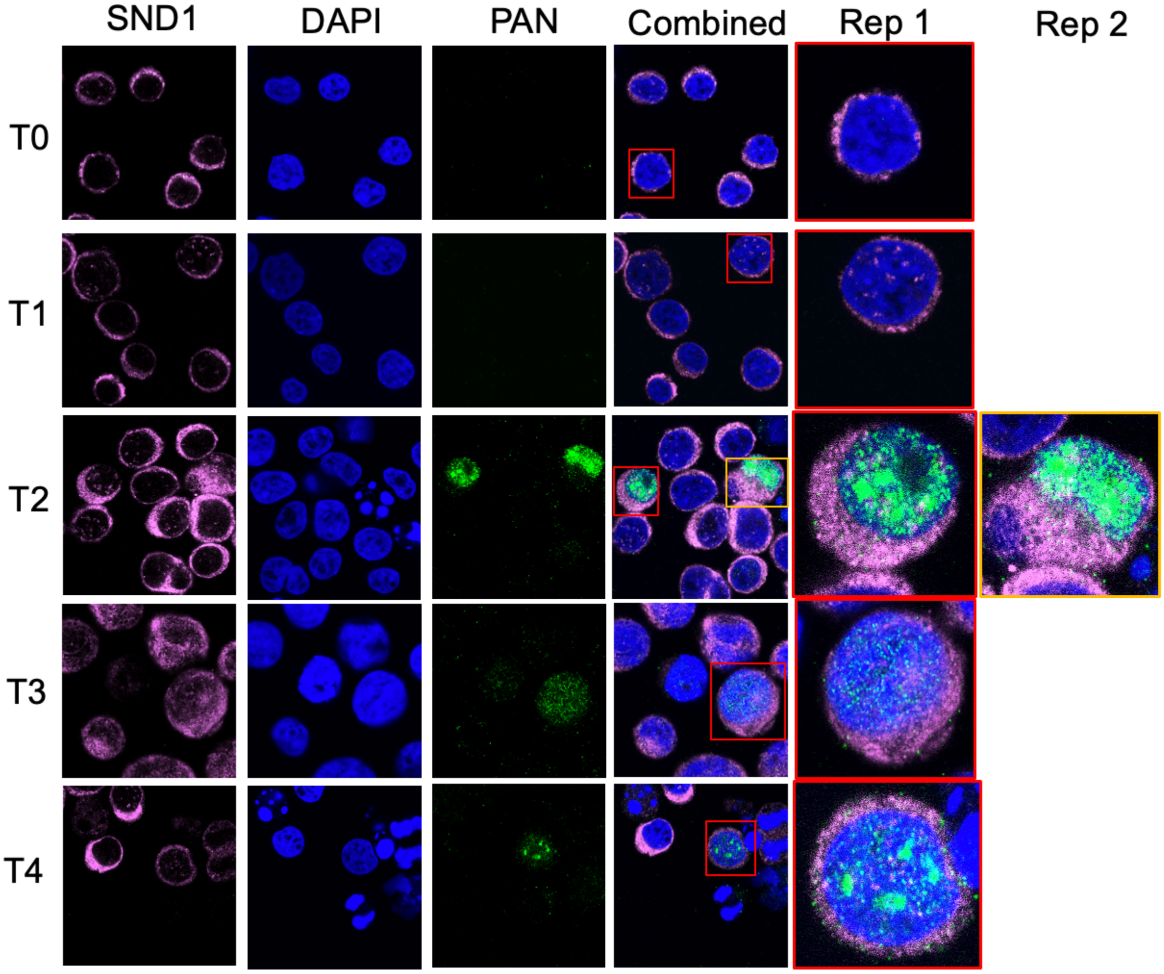

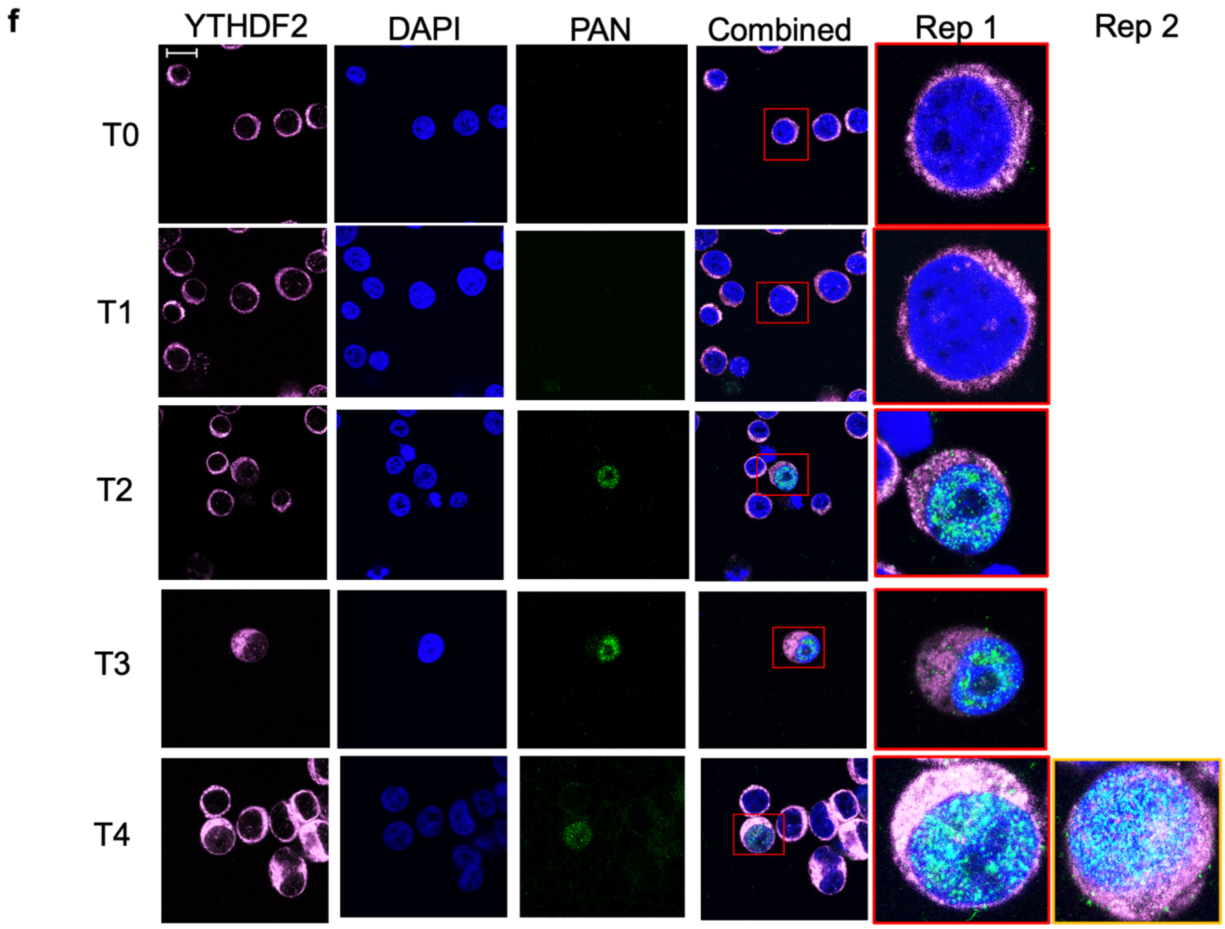


**Supplementary Fig. 10** Confocal fluorescent microscopy analysis of PAN and related m$^6$A methylome components performed in BCBL-1 cells during the latent (T0) and lytic (T1 – T4) stages of KSHV replication. The confocal microscopy micrographs representing the individual methylome enzymes (in pink) are ordered as follows: **a**, RBM15, **b**, METTL3, **c**, FTO, **d**, HNRNPC, **e**, SND1, and **f**, YTHDF2. DAPI staining of the nuclei is in blue, and PAN RNA is green.
